# Multiple roads to swarming: divergent molecular machineries drive the repeated evolution of locusts

**DOI:** 10.64898/2026.08.19.745877

**Authors:** Maéva A. Techer, Sheina B. Sim, Olga Dudchenko, Anna K. Childers, Julianne Allred, Emily Baker, David Mario Bellini, Devon Boland, Herman A. Dierick, Richard B. Dewell, Bert Foquet, Scott M. Geib, Ruqayya Khan, Richelle Marquess, Audélia M. C. Mechti, Martina E. Pocco, Kathryn Puperi, Seema Rana, Stephen Richards, Brian Scheffler, Tyler J. Simmonds, Amanda R. Stahlke, David Weisz, Fabrizio Gabbiani, Gregory A. Sword, Erez Lieberman Aiden, Hojun Song

## Abstract

Locust swarming, one of nature’s most spectacular examples of a repeated emergent polyphenism, has long been suspected to rely on conserved “swarming genes” or shared genomic features. By applying a model-clade approach comparing six species that vary in their degrees of plasticity and collective behavior, we show that the evolution of swarming locusts is not driven by shared genomic features or a universal set of swarming genes. In contrast, we find that this phenomenon evolved through flexible regulatory architectures, in which the degree of behavioral plasticity directly correlates with the total scale of density-responsive gene expression. While different locust species recruit largely non-overlapping gene sets to achieve the same syndrome, these divergent molecular machineries converge on similar higher-level biological functions. Thus, multiple molecular pathways achieve locust swarming, challenging the preconceived notion about the genetic prerequisites to transition from solitary to collective states. Further, we establish that a complex syndrome such as locust swarming emerges through modular regulatory systems that can be amplified, modified, or attenuated across the tree of life.

**Teaser:** There is no single molecular blueprint for becoming a swarming locust.

## INTRODUCTION

Locust plagues are one of nature’s most terrifying spectacles of destruction through collective transformation (*1*). They endanger the livelihoods and food security of 1 in 10 people worldwide. Their impact is comparable to that of other natural disasters, such as hurricanes or earthquakes, causing immense destruction in a short time (*2*). Ironically, these devastating insects also offer a gateway for unraveling the dynamic interaction between nature and nurture (*3*). Locusts display a spectacular example of density-dependent phenotypic plasticity, enabling them to coordinate distinct behavioral, physiological, and morphological traits in response to changes in population density (*4–6*). In response to an increase in density, normally cryptic and solitary individuals (solitarious phase) transform into conspicuous, gregarious individuals (gregarious phase). The behavioral transformation can occur within hours (*7*, *8*) and is followed by the expression of density-specific physiological (*9*), morphological (*10*), and ecological traits (*11*, *12*), ultimately culminating in massive swarms that can migrate thousands of kilometers (*13*) while wreaking havoc on agriculture and human livelihoods on a continental scale. This phenomenon is known as locust phase polyphenism (*14*). Understanding its genomic basis could lead to solutions for controlling these pests.

All locusts are grasshoppers, but not all grasshoppers are locusts. Currently, fewer than 20 of the nearly 7,000 described species of grasshoppers (Orthoptera: Acrididae) are known to exhibit the hallmarks of swarming locusts (density-dependent phenotypic plasticity and collective behavior) (*15*, *16*). Intriguingly, locusts are a phylogenetically heterogeneous group, suggesting that the ability to exhibit these complex coordinated alternative phenotypes leading to devastating migratory swarms has independently evolved repeatedly (*4*, *17*). This repeated emergence of locust swarming raises fundamental questions about its underlying genomic architecture and whether locusts have convergently evolved genomic features that set them apart from regular grasshoppers.

The genus *Schistocerca* represents a remarkable natural experiment that captures the evolution of phenotypic plasticity, adaptive radiation, and convergence. It contains about 50 species, of which three are swarming locusts and the remaining are non-swarming, sedentary grasshoppers (*4*). This genus also shows an unusual transatlantic disjunction, with only one species in Africa and the rest found throughout the Americas (Fig. 1A). The sole Old World representative is the Desert locust (*S. gregaria*), with historic plagues recorded in the Bible and Quran and continued plagues causing humanitarian crises, with the most recent being the East African plague (2019–2022). In the New World, the Central American locust (*S. piceifrons*), known as “saak” in Mayan culture, regularly affects a wide swath of tropical countries from Mexico to Panama (*18*) and has the potential to invade the United States as a changing climate may have altered the potential species range (*19*). The South American locust (*S. cancellata*) experienced a major outbreak in Argentina in 2015 after a 60-year period of recession, with swarms spilling over into neighboring countries (*20*). These locusts occupy vastly different habitats, from semi-deserts and deserts (*S. cancellata*, *S. gregaria*) to dry tropical woodland (*S. piceifrons*) and croplands during outbreaks (*21*). Previous phylogenetic studies strongly suggest that the ancestral *Schistocerca* originated in Africa and colonized the New World by transatlantic migration, implying that this ancestor must have been a swarming locust (*4*). As lineages diversified, descendant species lost their ability to swarm and became sedentary. Yet this ability re-evolved independently two separate times, in both *S. piceifron*s and *S. cancellata*, as they do not form a monophyletic group. Intriguingly, locusts are closely related to non-swarming, sedentary grasshoppers that can exhibit varying degrees of plasticity when experimentally induced. For example, the American bird grasshopper (*S. americana*) and the Cuban bird grasshopper (*S. serialis cubense*) form a clade with *S. piceifrons* and are known to show variable degrees of plasticity in terms of behavior and coloration, reminiscent of swarming locusts (*22*, *23*). Other sedentary species in the genus show much reduced or no plasticity, such as the gray bird grasshopper (*S. nitens*) (Fig. 1A). Therefore, the in-depth comparative study of the genomes of three locust species and three grasshopper species with varying degrees of plasticity and collective behavior within *Schistocerca* can reveal novel insights into the loss and gain of locust phase polyphenism in a phylogenetic framework.

**Figure 1:**
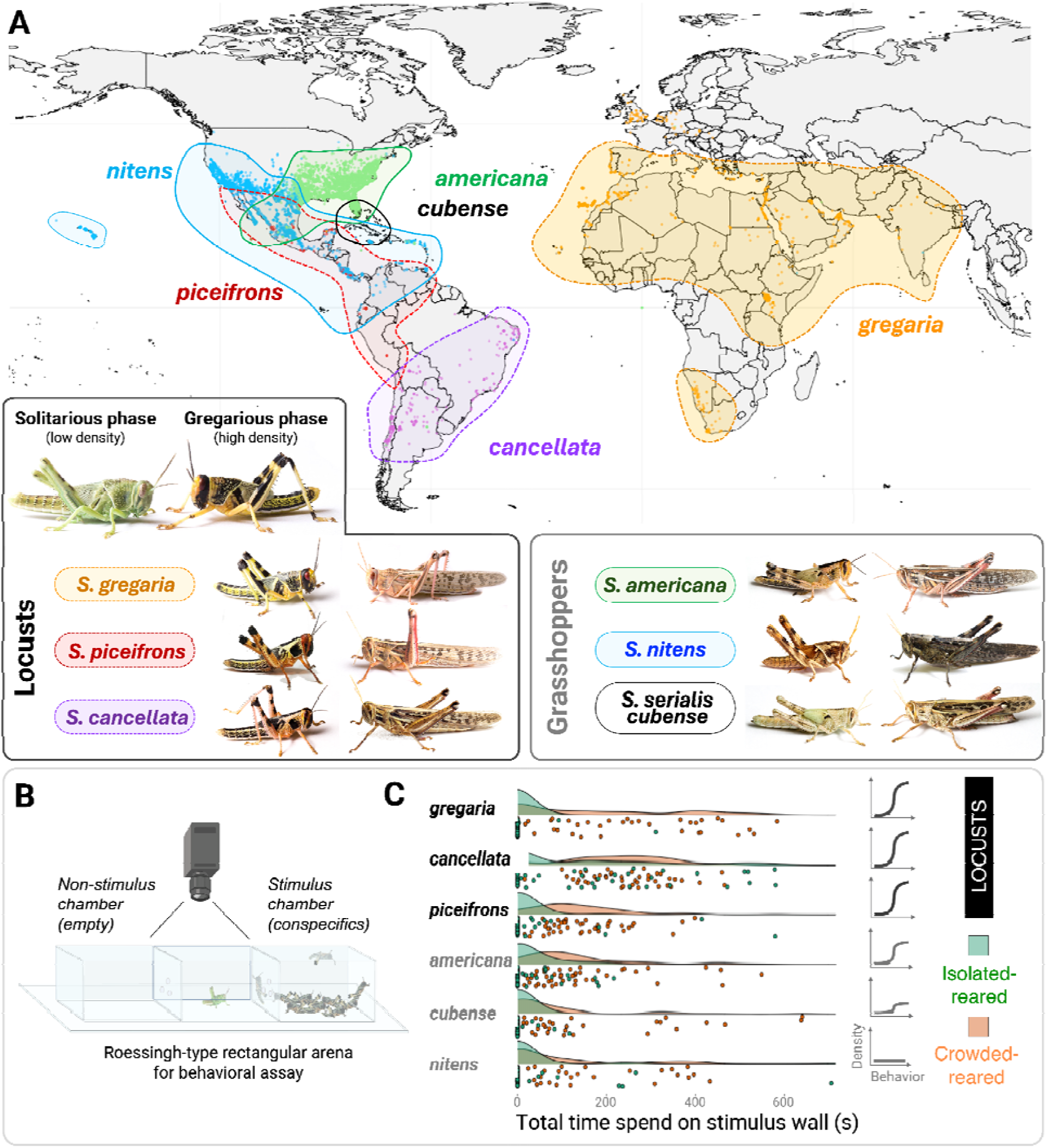
Density-dependent behavioral plasticity in six biogeographically distinct *Schistocerca* species. The six species compared include three swarming locusts and three non-swarming grasshoppers, each occupying different ecological and geographic ranges. These species exhibit a behavioral continuum in response to changes in density, which can be measured in a two-choice behavioral assay. Panel (A) depicts representative last-instar female nymphs and adults for all six targeted species, shown alongside, at the top, an example of *S. gregaria* in its solitary and gregarious phases for reference. The distribution map reflects biogeographic ranges from the literature (*2*, *20*, *33*, *123*, *124*), overlaid with species occurrences (dot symbols) from the GBIF database. Panel (B) illustrates the video-recorded behavioral arena design, where 50 conspecific juveniles serve as a social stimulus source, contrasted with non-stimulus control zones to quantify attraction and movement shifts. Panel (C) shows the total time spent on the stimulus wall by each species in a ten minute assay and summarizes overall behavioral reaction norms based on composite behavioral measurements, highlighting conserved density-dependent behavioral changes across the swarming species.

The study of locust phase polyphenism in such a comparative framework has been hampered by two primary challenges. First, locust and grasshopper genomes, ranging from 5 to 20 Gb (*24*), are among the largest within Insecta and contain complex repeats that are difficult to assemble and annotate (*25*). Only recently have sequencing technologies and bioinformatics tools become refined, integrated, and affordable enough to accurately generate such large genomes. Indeed, based on our survey of publicly available genome assemblies for the Orthoptera taxa in the NCBI, only five highly fragmented genomes were available prior to 2020. By early 2026, this number had increased to 81 genomes, of which 51 were chromosome-length. The second challenge is the difficulty associated with studying phenotypic plasticity across multiple species (*26*, *27*). Unlike most fixed traits that can be easily measured, phenotypically plastic traits must be revealed by exposing organisms to varying environmental conditions before quantifying the resulting reaction norms at the behavioral or gene expression levels (*28*). Generating such data under controlled conditions is especially challenging for non-model organisms, and even more so for multiple species on different continents. However, recent advances in standardized rearing and experimental manipulation of locusts, informed by comparative studies, are now making it possible to bridge these gaps and investigate phenotypic plasticity in a comparative molecular framework (*23*, *29*).

In this study, we addressed these challenges by combining single-molecule HiFi long-read sequencing with Hi-C short-read sequencing to assemble chromosome-level reference genomes of three swarming locust (*S. gregaria, S. piceifrons, S. cancellata*), and three non-swarming grasshopper (*S. americana, S. serialis cubense, S. nitens*) species of known phylogenetic relationships in the genus *Schistocerca.* We performed a comparative genomic analysis to test if there are specific genomic features that differentiate locusts from grasshoppers. We then quantified and compared density-dependent behavioral and transcriptomic reaction norms across all six species in a phylogenetic framework to test how plasticity evolved along the phylogeny. We identified candidate genes associated with locust phase polyphenism across the three locust species and tested their functions using RNA interference (RNAi) and behavioral assays. This revealed that locusts and grasshoppers differ primarily in their gene expression responses to crowding, despite having similar genomic backgrounds. We also show that different locust species evolved different molecular machineries to achieve similar swarming behaviors, representing an intriguing example of convergent evolution.

## RESULTS

### A spectrum of density-dependent reaction norms across *Schistocerca*

While locusts and grasshoppers are commonly distinguished by their collective behavior, a more critical distinction comes from the ability to express different coordinated phenotypes in response to changes in density. Previous studies have hinted that some non-swarming grasshopper species in *Schistocerca* carry hidden plastic reaction norms that can be experimentally induced (*22*, *23*, *30*, *31*), suggesting that density-dependent phenotypic plasticity could be a phylogenetically conserved feature within the genus, and that as a result *Schistocerca* locusts have independently evolved collective behavior. To examine this hypothesis and understand the genomic basis of phenotypic plasticity, we reared three swarming locusts and three non-swarming grasshopper species in both isolated and crowded conditions to induce density-dependent reaction norms (*29*), and quantified their resulting behaviors using a standardized locust behavioral assay (Fig. 1B and Supplementary Figure S1) (*32*). We combined previously published behavioral data (*23*) with newly collected trials to measure both activity-related and conspecific attraction-related parameters (Fig. 1C). Because data were collected in a standardized manner, we were able to make direct comparisons across the six species.

As expected, the three locust species showed highly plastic behavioral reaction norms in response to a change in density. Isolated nymphs showed low movement and strong avoidance of conspecifics, while crowded nymphs expressed high activity (distance moved, time spent moving) and attraction to conspecifics (positions in zones) (Supplementary Figure S1). In particular, the *S. gregaria* retained strong behavioral plasticity, with responses comparable to those of other locust species. Strikingly, *S. americana* exhibited plastic behavioral reaction norms indistinguishable from those of locusts, despite not swarming in nature. *S. serialis cubense* also showed plastic behavioral reaction norms, but at a reduced level, while *S. nitens* showed largely non-plastic behavioral reaction norms. The clade comprising *S. piceifrons*, *S. americana*, and *S. serialis cubense* is particularly intriguing because these species exhibit the clearest spectrum of plasticity and can help illuminate how locust phase polyphenism might have evolved. It is conceivable that the common ancestor of these three species could have experienced heterogeneous environmental conditions, which may have driven the evolution of greater plasticity. However, the degrees of plasticity differ among these species, suggesting that either the evolution of plasticity within the clade was gradual or that what we observe is the end result of selection or drift shaping the degree of plasticity (*28*). Thus, comparing locust and grasshopper species may help reveal genomic features associated with varying degrees of plasticity and collective behavior.

### Chromosome-length genome assemblies reveal a conserved genomic architecture across locusts and grasshoppers

We generated chromosome-level reference genomes for our six target species that vary in their degrees of density-dependent phenotypic plasticity (Fig. 1C). For *S. gregaria*, one of the most well-studied locust species, we generated a high-quality reference genome by combining PacBio HiFi long reads with Hi-C-linked Illumina short reads. Using the same methodology, we also generated reference genomes of the other two *Schistocerca* locusts: *S. piceifrons* and *S. cancellata,* and for three non-swarming grasshopper target species in the genus, giving genomes of similar contiguity and accuracy. This allowed us to test whether genomic features associated with locust swarming exist and if they are phylogenetically conserved or divergent.

The six datasets spanned 8.6-9.1 Gb (Fig. 2A and Supplementary Table S1), placing them among the largest insect genomes assembled at chromosome-length. We found that genome sizes differed only modestly between locusts (8.67 ± 0.12 Gb) and grasshoppers (8.97 ± 0.13 Gb). For *S. serialis cubense*, the genome size was 9.1 Gb, making it the largest insect reference genome currently available. Scaffolds were highly contiguous (Supplementary Table S1) and assemblies show uniformly high completeness, with BUSCO scores exceeding 99%, comparable to other smaller-sized insect reference genomes (Fig. 2A). *S. gregaria* not only was more contiguous than the previous draft genome assembly (*33*) but also had better contiguity (lower scaffold number, and contig N50) than more recent independent locust assemblies for *S. gregaria* and *L. migratoria* (*34*) and genome completeness (Table 1).

**Figure 2:**
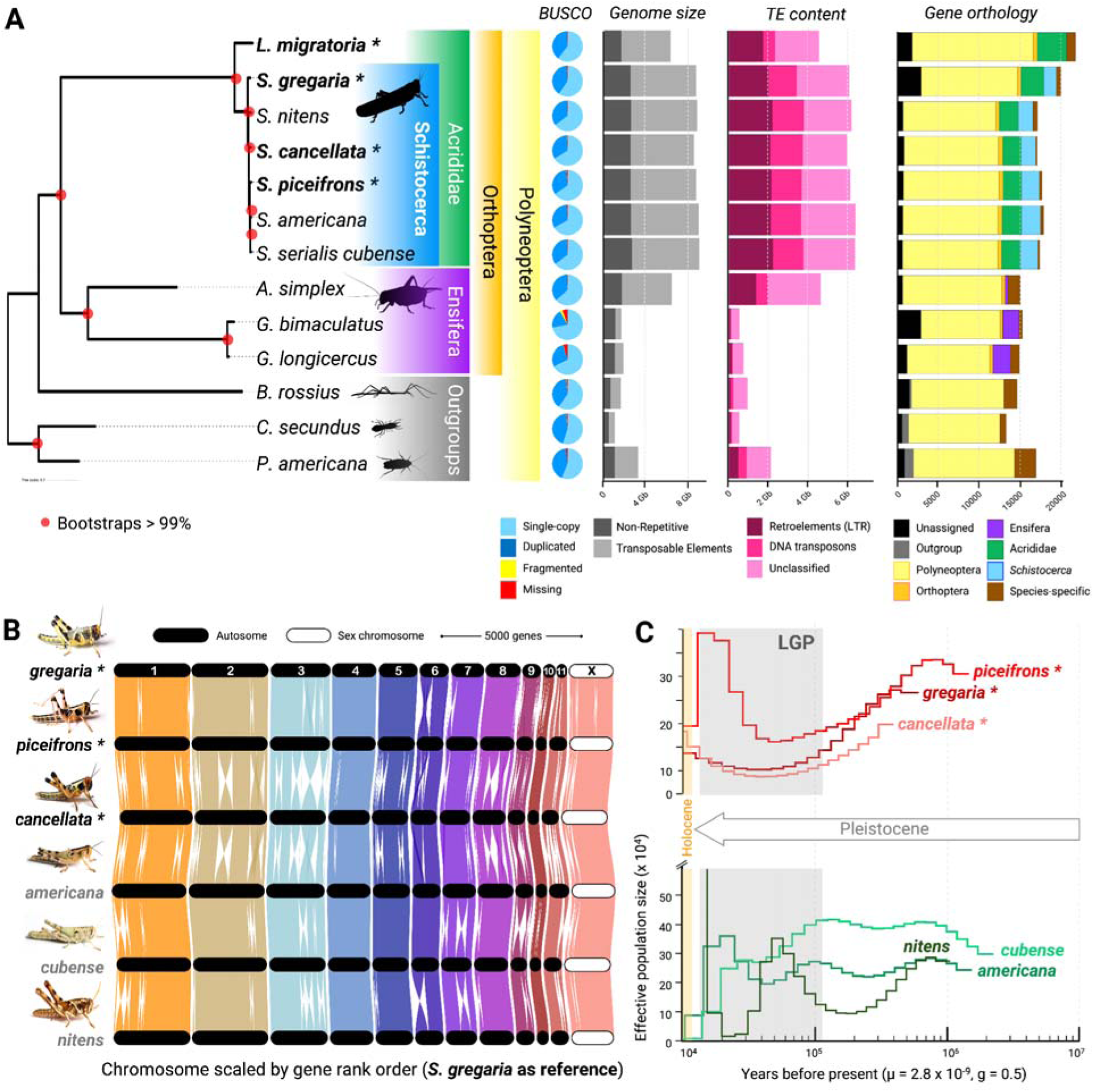
*Schistocerca* genomes provide a comparative framework to investigate genomic features shared by locusts. (A) Phylogenetic placement of the three *Schistocerca* **locusts** (*) and three grasshoppers sequenced (boxed in blue), alongside other Orthoptera and Polyneoptera outgroups, based on 3,128 single-copy orthologs. For each genome, BUSCO scores (genome completeness), genome size, and transposable elements (TE) content and composition are shown. The gene orthology profiles, color-coded by the phylogenetic lineage, highlight the proportion of shared and lineage-specific genes per species. (B) Whole-genome synteny plots demonstrate highly comparable chromosome-scale architectures and collinearity within the genus using *S. gregaria* as a reference, with conserved gene content and large inversion in some autosomes. (C) Single-genome pairwise sequentially Markovian coalescent (PSMC) reconstructions reveal contrasting demographic histories between locusts and grasshoppers *Schistocerca*, with each line representing effective population-size variation (Ne) from 100 bootstraps for each species and considering 2 generations per year. The gray shaded area shows the last glacial period (LPG).

**Table 1:**
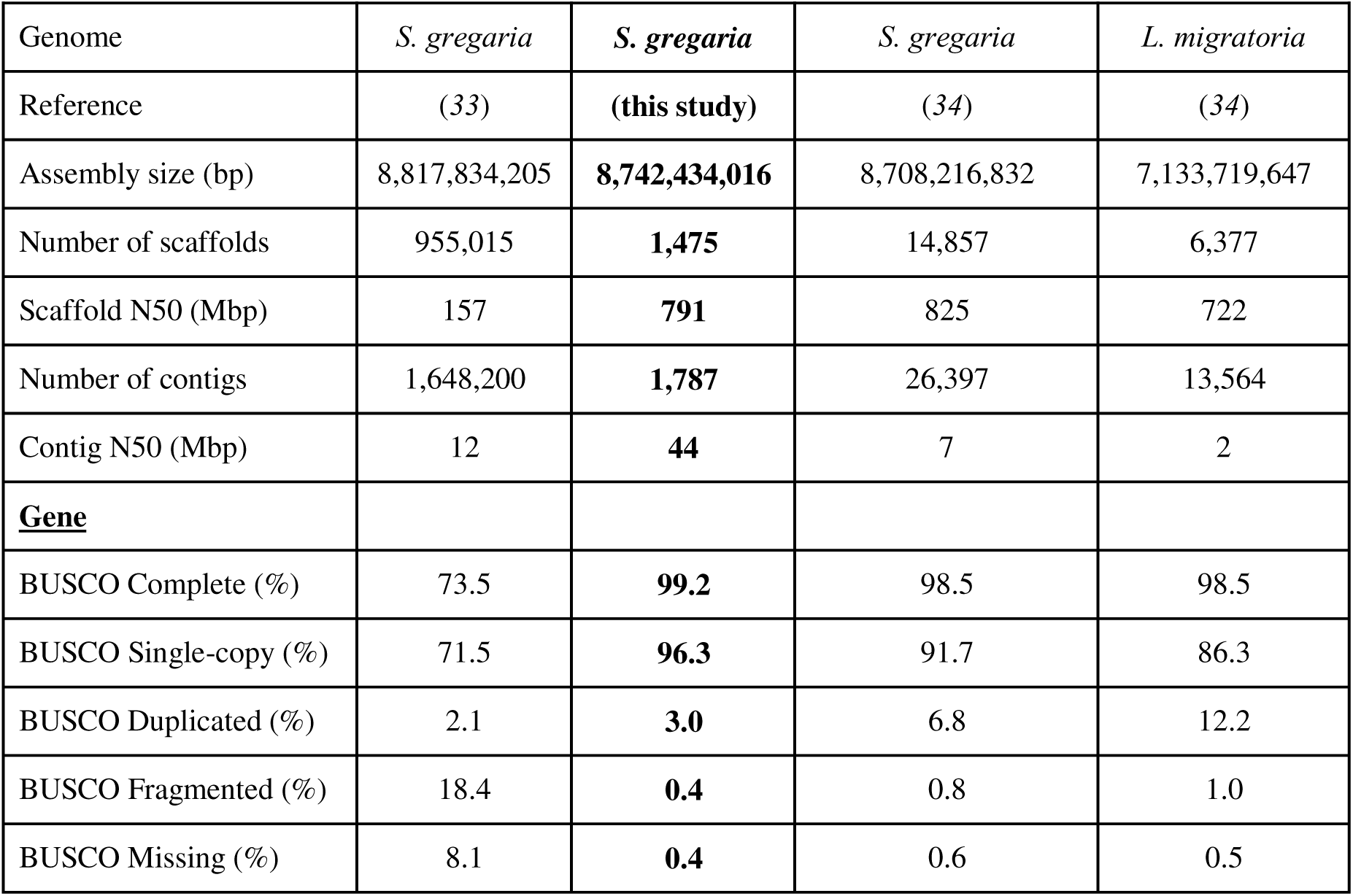
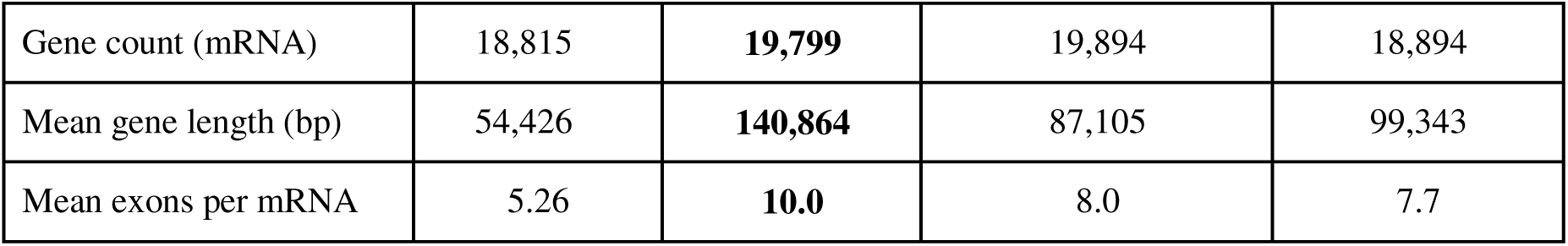
Comparative summary statistics on genome contiguity and gene content for foundation model species in locust phase polyphenism: the desert locust *S. gregaria* and migratory locust *L. migratoria* assemblies. In bold, are highlighted the statistics of our assembly which is the best among our six *Schistocerca* reference genomes.

Despite their extremely large sizes, all assemblies recovered the canonical cytogenetic karyotype in *Schistocerca* of 11 autosomes plus an X chromosome (Fig. 2B and Supplementary Figure S2) (*35*, *36*). The largest autosome corresponding to L1 in (*35*), contributed 1.20 to 1.32 Gb per assembly, larger than most complete insect genomes. Whole-genome alignments (NCBI Comparative Genome Viewer) and ortholog-based synteny analyses (GENESPACE) revealed extensive conservation of gene order across chromosomes, with only modest large-scale rearrangements (Fig. 2B). The fourth autosome was the most collinear chromosome across all species, whereas other chromosomes carried species-specific structural polymorphisms. Thus, at the gross level of chromosomal structure, gene content, and syntenic patterns, the genomes of locusts and grasshoppers were highly conserved and did not show any specific differences associated with swarming, prompting us to examine other genomic features.

### Species-specific and large repeat contents characterize locust and grasshopper genomes

Large eukaryotic genomes are often highly repetitive (*37–39*). We found that this was true for *Schistocerca* genomes, as they contained 70.6-71.7% repeat sequences (Fig. 2A), placing them among the most repeat-rich insect genomes to date (*40*). We annotated a subset of repeats as transposable elements (TEs), although most repetitive regions remain unclassified. DNA transposons, such as long interspersed elements (LINEs), long terminal repeats (LTRs), and unclassified elements, occur in broadly similar proportions across species (Supplementary Data S1). Although the functional roles of TEs in *Schistocerca* remain unclear, studies in other systems identify TEs as major sources of genetic and regulatory variation and as potential modulators of gene expression and phenotypic plasticity (*41*).

Given the relatively young age of the *Schistocerca* genus (∼7.9 million years (*4*)) and its conserved chromosomal architecture, we expected repeat landscapes to be similar. Instead, Kimura distance–based repeat age distributions (reflecting repeat age-structure) revealed striking interspecific differences (Supplementary Figure S3). The repeat landscape of *S. gregaria* was characterized by several major and minor bursts, all relatively recent (0-17% divergence). Although recurrent boom-bust population cycles might facilitate dynamic TE accumulation, the relationship between effective population size and TE content remains unresolved (*42*). However, the repeat landscapes of the other two swarming locust species do not resemble that of *S. gregaria*, indicating that TE dynamics differ even among swarming species. Instead, closely related species show more similar repeat age structures. *S. piceifrons*, *S. americana*, and *S. serialis cubense,* which form a monophyletic clade, share broadly comparable landscape profiles characterized by two major bursts, although the detailed signatures differ. *S. piceifrons* showed a prominent intermediate-divergence wave of LINEs and transposons, while the other two species retained signatures of deeper and older TE proliferation waves (Supplementary Figure S3). Because these species diverged less than ∼2 million years ago (*4*), repeat landscape shape remained conserved over short evolutionary timescales while shifting along lineage-specific trajectories. Interestingly, the swarming *S. cancellata* and non-swarming *S. nitens* display similar repeat landscapes despite their distant relationship. Collectively, these patterns indicate that TE accumulation follows species-specific evolutionary trajectories shaped by shared ancestry rather than by swarming behavior.

### Locust and grasshopper demographic histories show contrasting signatures consistent with differences in population dynamics

We reconstructed changes in effective population size (Ne), as a genetic proxy for long-term population size, using single-genome Pairwise Sequentially Markovian Coalescent (PSMC) analyses (Fig. 2C). As grasshopper and locust ecology can vary with resource availability, climatic conditions and phase, we assessed the robustness of inferred Ne trajectories by modulating generation time (from univoltine to trivoltine (*21*)) (Supplementary Figure S4). Overall, inferred demographic trajectories were robust to generation time assumptions, and we therefore used an average of two generations per year for cross-species comparability.

Sedentary, non-swarming species show gradual expansions and contractions in Ne that broadly track major climatic oscillations, including glaciation and associated vegetation shifts across the Americas (*43*, *44*). These trajectories resemble demographic patterns of many temperate organisms responding to Quaternary climate change (*45–47*), with range expansions during favorable periods and contractions during colder or drier phases. In contrast, swarming locust species do not exhibit the same clear climatic imprint, with a smoother Ne recovery. Their demographic trajectories appear more diffuse and less tightly coupled to glacial cycles. Recurrent transitions between the large, panmictic gregarious phase and the localized, solitarious phase likely repeatedly restructured genetic variation. This pattern suggests that the unique population dynamics of swarming cycles followed by a bottleneck can obscure long-term climatic signals in locust genomes compared to grasshoppers. While this may limit the resolution of ancient demographic inference with a single genome, it highlights a key biological insight: swarming behavior itself reshapes population genetic history, decoupling it from long-term environmental fluctuations.

### Lineage-specific gene family expansions are associated with partially shared functional pathways in locusts

Aside from the repeats, large genomes are often characterized by increased gene contents (*48*). We explored whether this pattern holds for *Schistocerca* and whether specific expansions are associated with locust swarming. The annotation of *S. gregaria* was supported not only by 1.7 billion short reads of RNAseq data but also by extensive IsoSeq sequencing (∼5 million reads), and 1.35 million transcriptome shotgun assemblies (TSAs), which yielded the most comprehensive annotation. This resulted in a total of 19,799 low-fragmented genes predicted in *S. gregaria* (Table 1). In comparison, genome annotation using the NCBI Eukaryotic Genome Annotation Pipeline (EGAP) predicted ∼17,100–17,700 protein-coding genes for our other *Schistocerca* genomes, which had less transcript evidence available to support annotation. Consistent with this difference, the proportion of gene models fully supported by transcript evidence was higher in *S. gregaria* (∼82%) than in the other species (∼64 - 71%). The inclusion of IsoSeq data further improved the resolution of full-length transcripts and alternative isoforms in *S. gregaria*. Despite their massive genome sizes, gene density was similar across species and remained relatively low (2.26 genes/Mb in *S. gregaria*; 1.99 ± 0.01 in the other locusts; 1.95 ± 0.04 in grasshoppers) when compared to the migratory locust *Locusta migratoria* (3.42 genes/Mb) and other Polyneoptera (a group of closely related insect orders including Orthoptera, Phasmatodea, and Blattodea). Across the six *Schistocerca* species, > 94% of the 106,025 genes cluster into 16,681 orthogroups (i.e., gene families), confirming the strong correspondence among gene families (Fig. 2A and Supplementary Data S1). Among these groups of genes, only 3.7% were species-specific orthogroups, and 52.6% consisted of single-copy orthologs (Supplementary Data S2). Most species shared >75% 1:1 orthologs with *S. gregaria*, although *S. piceifrons* showed a lower proportion (67.0%) and more 1:N relationships, suggesting lineage-specific duplications or annotation differences. Gene annotations and orthology inference provided a framework to investigate lineage-specific expansions and contractions.

Using OrthoFinder (*49*) and CAFE5 (*50*), we inferred gene family expansions and contractions across Polyneoptera (Fig. 3A and Supplementary Data S3). The genus *Schistocerca* shows substantial ancestral gene family gains and a relatively balanced gain-loss ratio, except for *S. nitens*, which had the highest number of net gene losses (-498 genes). Swarming locusts show expansions in gene families associated with energy metabolism, including those involved in long-distance flight. *S. gregaria* exhibits expansion of NADH–ubiquinone oxidoreductase components (+77 genes in OG0000041), consistent with previous findings in *L. migratoria* (*51*). *S. cancellata* shows expansion in gene families linked to craniofacial development (+13 genes in OG0000024), and *S. piceifrons* expands the DNA repair protein RAD50 family (+37 genes in OG0000150). Histone gene families expand extensively in *L. migratoria* (+102 genes across multiple orthogroups). When looking at the identity of the expanding and contracting families, we found no complete overlap or candidate gene families shared by all locusts (Fig. 3B). Instead, Kyoto Encyclopedia of Genes and Genomes (KEGG) enrichment analyses identified pathways involved in DNA folding, sorting, degradation, and neurodegenerative processes among gene-gain categories shared by the three *Schistocerca* locust species (Fig. 3C).

**Figure 3:**
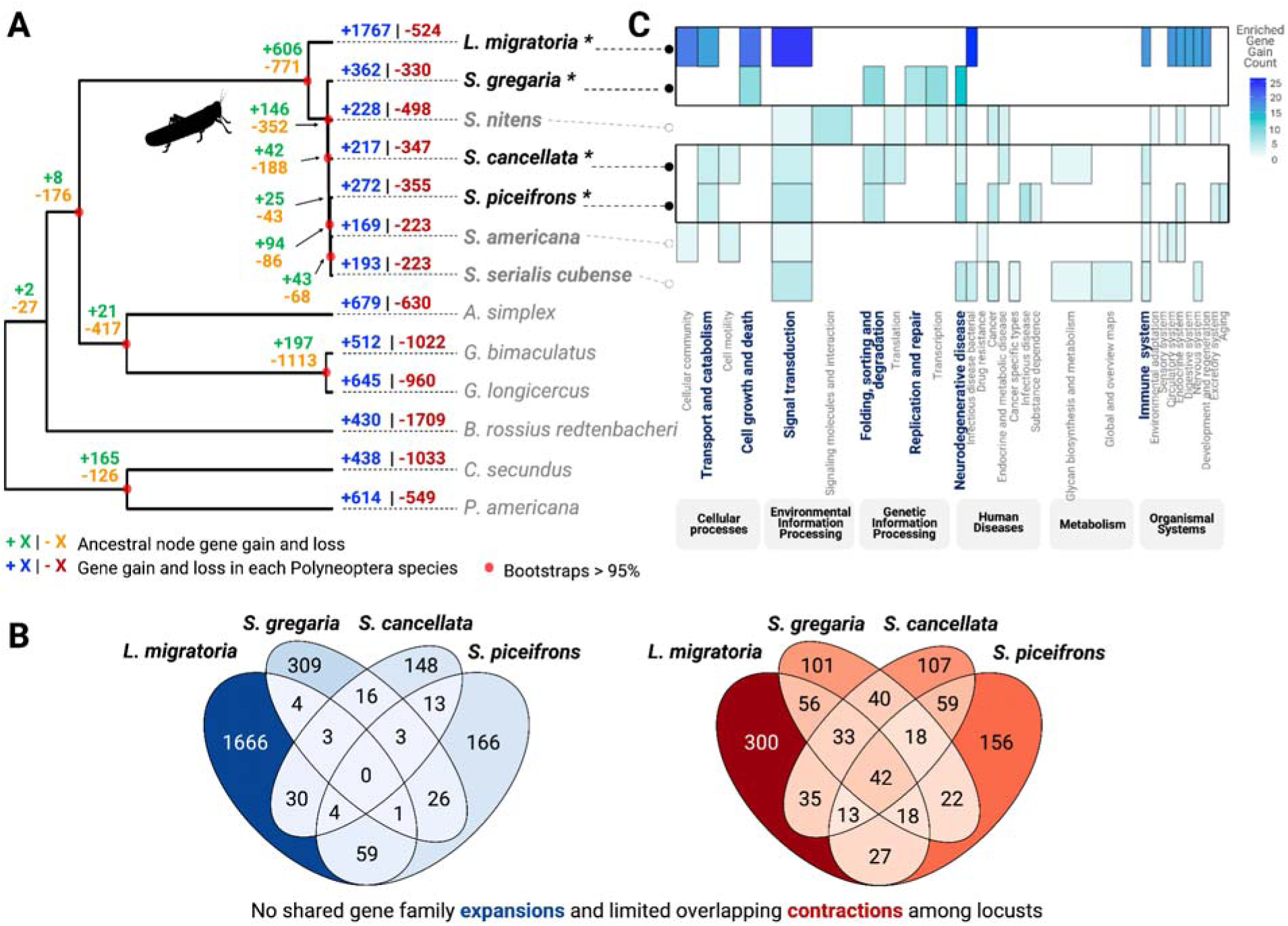
Swarming and non-swarming *Schistocerca* species show subtle but divergent patterns of gene famil evolution associated with the swarming trait. (A) Gene family expansion (positive values) and contractions (negative values) across *Schistocerca* and Polyneoptera outgroups. (B) While overall rates of losses and gains were similar within *Schistocerca*, except for *S. nitens*, the pathways of the gene families involved were markedly different among species. (C) Venn diagrams of shared orthogroups/gene families found expanding or contracting across four locust species.

We found a signal of shared gene families lost among locusts, but it appears that this signal was associated with high levels of expansions in sedentary species (Fig. 3B). Indeed, non-swarming grasshopper species harbored multiple large species-specific expansions in orthogroups with broad or unknown functions (*S. americana*: +17 genes in OG0000145, *S. serialis cubense*: +28 genes in OG0000078, *S. nitens*: +30 genes in OG0000331) and in gene families not clearly linked to behavioral plasticity, such as collagenase 3 and matrix metalloproteinase (*S. nitens*: +21 genes in OG0000670) (Supplementary Data 3 and 4). Taken together, gene family evolution reveals lineage-specific expansions rather than a single convergent set of “swarming genes.” Swarming locusts do not share a uniform gene family expansion signature; instead, each species exhibits distinct molecular elaborations consistent with independent evolutionary trajectories.

### Signals of episodic positive selection follow lineage-specific trajectories rather than a shared swarming signature

Given that locust phase polyphenism, coupled with collective behavior, is considered an adaptation to heterogeneous environmental conditions (*52*), we explored whether certain genes exhibit patterns consistent with positive selection. We tested for signatures of selection using single-copy orthologs at two phylogenetic scales (Polyneoptera-wide and Acrididae-specific), applying an adaptive branch-site random effects likelihood model (aBSREL) and the Branch-Site Unrestricted Statistical Test for Episodic Diversification for phenotypes, as well as relaxed selection (implemented in BUSTED-PHenotype + RELAX) to test for positive, episodic, and relaxed selection associated with swarming lineages (*53–55*) (Fig. 4A). Across the 761 1:1 Acrididae orthogroups, 6.8% of branches tested showed significant evidence of positive selection at one or more sites using aBSREL (Supplementary Data S4). *L. migratoria* carried the highest number of selected branches (n = 94), followed by other locusts (*S. piceifrons* n = 73; *S. cancellata* n = 63; *S. gregaria* n = 51) and grasshoppers (*S. americana* n = 50; *S. nitens* n = 44; *S. serialis cubense* n = 41). Analyses of Polyneoptera-wide 1:1 orthologs produced comparable patterns (Supplementary Figure S5). When considering all 1:1 ortholog lineage classes (Fig. 1A), we found a reduced overlap with only two orthogroups for which all locusts show positive selection (Fig. 4B). These genes were Acrididae-specific, annotated as carboxypeptidase N subunit 2-like or uncharacterized. We also did not detect a signal of selection in senataxin (*Setx*) genes, previously hypothesized to be associated with the swarming phenotype (*56*).

**Figure 4:**
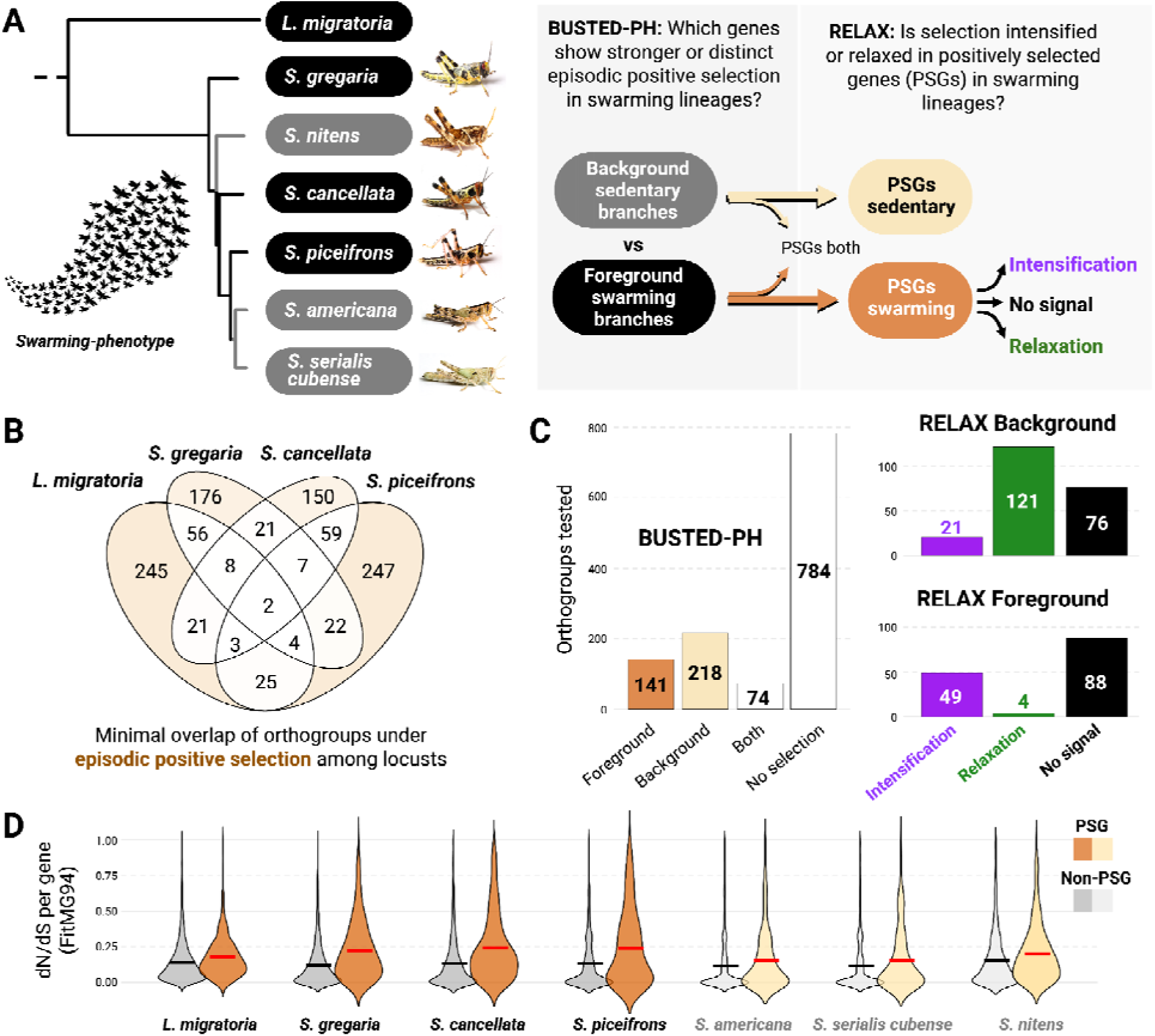
Episodic positive selection is widespread but low-level, with divergent patterns, limited overlap, and signals of intensification in swarming locusts. (A) Overview of the selection analysis workflow and pruned phylogeny used for the phenotype contrasting BUSTED-PH and RELAX analysis across the Acrididae. (B) Venn diagram presenting the minimal overlap of orthogroups for which locust species showed branch level signal of episodic positive selection with aBSREL. (C) Genome-wide selection patterns for PSGs (Positively Selected Genes detected by BUSTED-PH) indicating the frequency of positive selection in sedentary versus swarming lineages and the intensity of the selection signal with RELAX along the branches tested. (D) Distribution of dN/dS ratios per PSGs versus non-PSGs uncovers that some locusts, such as *S. piceifrons* and *S. cancellata,* exhibit the strongest signatures of diversifying and episodic selection across their genomes.

However, genome-wide tests on the Acrididae phylogeny using BUSTED-PH to contrast selection across phenotypes (Fig. 4A) revealed that genes without detectable selection dominated the distribution (χ² = 1042.8, df = 3, p < 0.05). Contrary to expectations, we found that more genes experienced selection on non-swarming (“background”) branches than on swarming (“foreground”) branches (Fig. 4C). However, direct comparisons indicated significantly stronger selection signals in background branches (χ² = 16.5, df = 1, p < 0.05) in both full and Acrididae-pruned phylogenies. Although swarming species did not show an excess of positively selected genes (PSGs) relative to grasshoppers, we identified 141 genes under selection in at least one locust lineage and not under selection in the sedentary lineages (n = 5,339 orthogroups tested) (Fig. 4C, Supplementary Data S5). Among these, *S. piceifrons* and *S. cancellata* exhibited the highest non-synonymous to synonymous substitution ratios (dN/dS) (Fig. 4D). Interestingly, PSGs along sedentary species branches showed more signals of relaxed selection, whereas locust lineages more frequently showed signals of selection intensification (Fig. 4C). This asymmetry in selection regimes between sedentary and swarming lineages highlights differences in how evolutionary pressures act on their genomes, despite the absence of a shared set of positively selected genes. Together, these results indicate that positive selection acts on distinct gene subsets across lineages rather than on a shared set of “swarming genes”. This suggests that swarming behavior does not correspond to a unified molecular signature of adaptive evolution but instead arises from partially overlapping, lineage-specific trajectories.

### Degree of plasticity across species correlates with the number of differentially expressed genes

Behavioral assays revealed a spectrum of density-dependent plasticity across species (Fig. 1C), but genomic complexity did not correlate with behavioral variation (Figs. 2-4). We therefore tested whether differences in density-responsive gene expression explain interspecific variation in plasticity and collective behavior. We induced density-dependent reaction norms in the laboratory and generated transcriptomes of head and thorax tissues from both isolated and crowded nymphs across all six species (Fig. 5A). We hypothesized that if gene expression patterns could explain plasticity, there would be numerous density responsive differentially expressed genes (DR-DEGs) between isolated and crowded individuals of the species showing behavioral plasticity, while there would be little or no DR-DEGs in the species showing reduced or no plasticity. To ensure robust cross-species comparisons, we identified DR-DEGs using two complementary approaches: mapping reads to each species’ own genome and mapping all reads to the best-annotated reference genome (*S. gregaria*). Both approaches produced concordant results (Supplementary Data S6).

**Figure 5:**
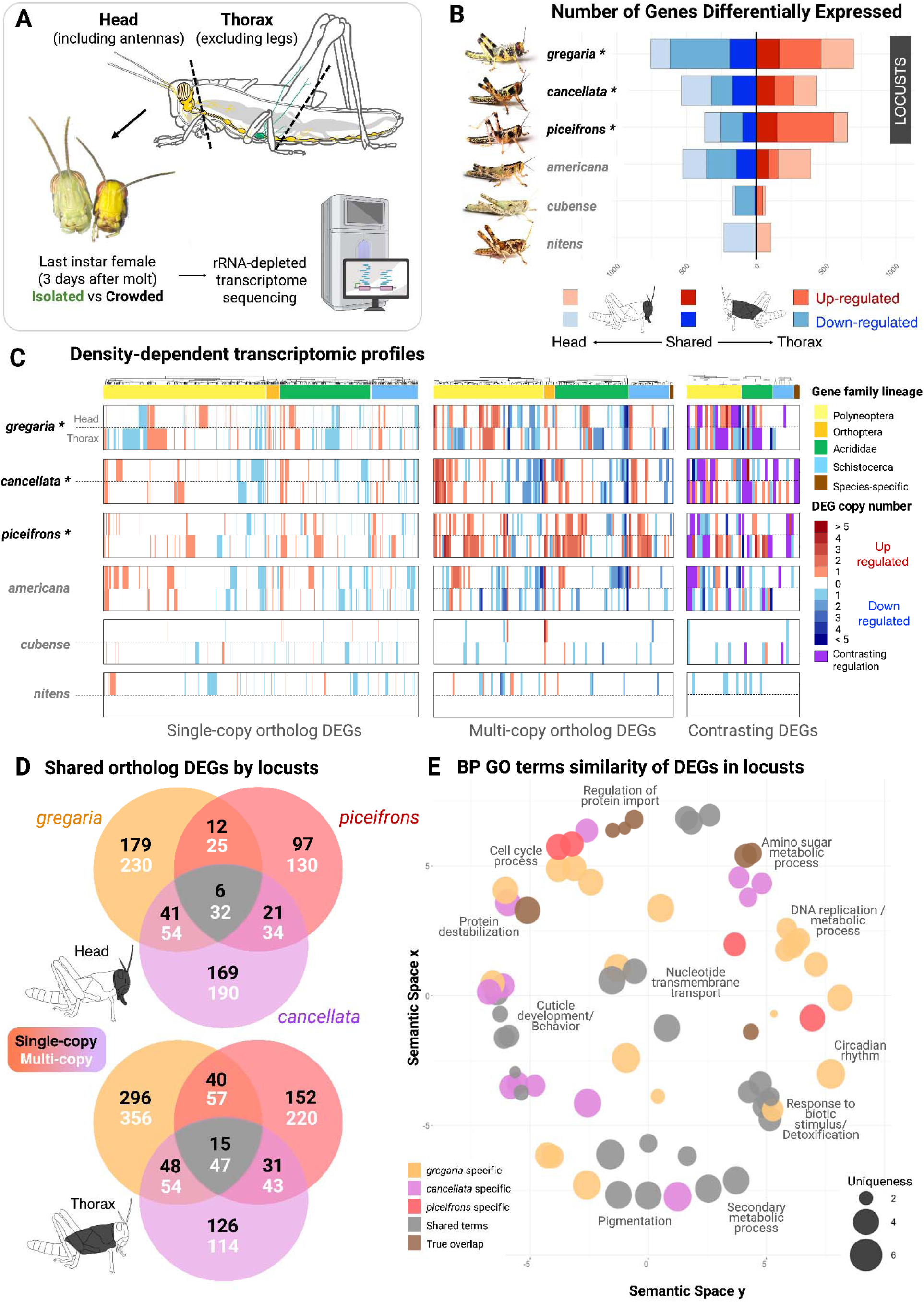
Density-dependent behavioral plasticity corresponds to heterogeneous transcriptomic responses across species. (A) The tissues targeted for density-responsive transcriptomes comparison were the whole head and thorax of last instar female nymphs, 3 days after molting. The anatomical side view shows the brain tissues in yellow and green targeted by the dissection of head and thorax. (B) Locusts (*) and *S. americana* showed the largest numbers of density-dependent differentially expressed genes (DR-DEGs) between isolated- and crowded-reared conditions in head and thorax tissues. Bars show upregulated (red) and downregulated (blue) DR-DEGs, with “Shared” indicating overlap between tissues.(C) Heatmaps of single-copy ortholog, multi-copy ortholog, and DR-DEGs with contrasting regulation show that each species and tissue displays a unique density-dependent transcriptomic profile. Columns are color-coded by gene family lineage, and rows indicate tissues. The DR-DEG copy number is shown by color intensity. (D) Although responses are vastly different, an overlap in DR-DEGs exists across the three swarming locusts. Venn diagrams show the overlap of single-copy (black label) and multi-copy (white label) DR-DEGs for each tissue. (E) Semantic similarity clustering of biological process (BP) gene ontology (GO) terms enriched among DR-DEGs in swarming locusts. Each circle represents a GO term, sized by the number of genes, with color indicating whether it is species-specific or shared.

As expected, all three locust species exhibited large numbers of DR-DEGs, in the hundreds per tissue and regulation direction (up-or downregulated) (Fig. 5B). *S. gregaria* showed the highest DR-DEG counts, followed by *S. piceifrons* and *S. cancellata*. DR-DEG numbers differed between tissues: the thorax contained more DR-DEGs than the head in *S. gregaria* (n = 1,083) and *S. piceifrons* (n = 812), whereas *S. cancellata* showed more DR-DEGs in the head (n = 679) (Supplementary Data S7). Among the non-swarming grasshopper species, *S. americana* exhibited the largest number of DR-DEGs, while both *S. serialis cubense* and *S. nitens* showed few or no DR-DEGs. As such, DR-DEG numbers scaled with behavioral plasticity across species, indicating an association between transcriptional responsiveness and reaction norm strength.

The three locusts and *S. americana* showed a large number of DR-DEGs (Fig. 5B), so we asked whether there exists a set of phylogenetically conserved “plasticity” genes; a group of genes that are DR-DEGs across species. However, orthogroup-resolved DR-DEGs showed that each plastic species used a distinct combination of density-responsive genes, including single-copy, multi-copy, and copy-variable families from various gene family lineages, rather than a phylogenetically conserved pattern (Fig. 5C and Supplementary Figure S6). Put simply, despite similar reaction norms in behavior and other phenotypes, the species regulated different genes in response to density. Among the three *Schistocerca* locust species, the overlap in DR-DEG identity remained low. Only 32 orthogroups in the head tissue and six orthogroups in the thorax were shared across the locusts. Shared single-copy genes were even fewer (six in the head and 15 in the thorax), representing a small fraction of each species’ total DR-DEG repertoire (Fig. 5D), even when compared to *L. migratoria* (Supplementary Figure S7). Shared orthogroups from the head tissue included multi-copy genes related to inositol oxygenase and peroxidase activity, cuticle structure and coloration, cytochrome P450, and ATP-binding cassette transporters, as well as single-copy genes associated with mucin, neuropeptides, and juvenile hormone signaling. Shared orthogroups from the thorax similarly involved mucin, cuticle proteins, juvenile hormone pathways, but also translation initiation factor 2F proteins. These overlaps indicate that certain functional modules recur across species, but the majority of DR-DEGs remain species-specific.

Comparisons between the sister species *S. piceifrons* and *S. americana* further illustrate this pattern. Both species exhibit strong behavioral plasticity and substantial numbers of DR-DEGs, yet their overlap is relatively low despite their close phylogenetic relationships. Moreover, the swarming *S. piceifrons* exhibited a larger DR-DEG repertoire across tissues than *S. americana*. The overall pattern strongly suggests that phenotypic plasticity across species, even within the same genus, is achieved through distinct molecular mechanisms, again indicating that there are several ways to evolve swarming locusts.

### Plasticity is driven by different density-responsive genes that are functionally similar

Although the identities of DR-DEGs differed across the three locust species, we asked whether they could serve similar functions. We conducted functional annotation of the DR-DEGs and performed enrichment analyses to identify overrepresented biological processes. Gene ontology (GO) analyses of biological process (BP) categories revealed significant enrichment for metabolism, neuromodulation, developmental regulation, cuticle remodeling, and stress-response pathways in the most behaviorally plastic species and across swarming locusts (Fig. 5E). Semantic similarity clustering of enriched GO terms showed a visible partial overlap of downstream processes among the three locust species, while also revealing species-specific enrichments. Similar patterns were observed for molecular functions, cellular components, and overall KEGG pathways (Supplementary Figure S8 and S9). Thus, although individual DR-DEGs differ substantially across species, the functional themes they represent remain similar. Distinct gene sets repeatedly target overlapping biological processes associated with metabolism, neural signaling, and morphological regulation. These results indicate that density-dependent behavioral plasticity converges at the level of functional pathways rather than at the level of gene identity.

### Co-expression networks underlying locust phase polyphenism are large and species-specific

Because we quantified both behavioral reaction norms and DR-DEGs within each species, we next tested whether co-expressed gene modules are associated with specific behavioral traits relevant for locust swarming. We constructed gene co-expression networks separately for each species and identified multiple modules of co-expressed genes. We filtered and explored the modules whose eigengene expression correlated significantly with behavioral parameters, including distance moved (activity) and time spent in the stimulus zone (attraction). In each *Schistocerca* locust species, we identified multiple modules significantly associated with these behavioral traits (individual module dendrograms as well as size- and module-behavioral trait relationships are available online (*57*)). Each module comprised hundreds to thousands of co-expressed genes, indicating that locust phase polyphenism is a complex, multigenic syndrome (Supplementary Figure S10). These patterns were consistent across tissues and species. In *S. gregaria*, multiple candidate genes clustered into two major modules, each exceeding 2,000 genes, and these modules, using the “unsigned” method, included hundreds of DR-DEGs that were jointly up- or down-regulated (Supplementary Figure S11). Thus, the transcriptional architecture underlying locust phase polyphenism appears distributed across extensive regulatory networks instead of a compact pathway.

Our comparative analyses further revealed species-specific organization of these networks. *S. gregaria* and *S. cancellata* displayed consistent module-trait correlations across behavioral variables, whereas *S. piceifrons* exhibited an inverse correlation pattern for time spent in the non-stimulus zone (Supplementary Figure S10). These contrasting network-trait relationships for density-dependent behaviors arise from differently structured gene co-expression architectures across species, bolstering the idea that divergent molecular machinery underlies similar behavioral responses to density.

### Phylogenetically conserved toolkit genes implicated in phenotypic plasticity are part of the genetic repertoire of locust phase polyphenism

While genomic and transcriptomic analyses indicate that different locust species have evolved distinct molecular machineries underlying locust phase polyphenism, we identified a small number of DR-DEGs shared among the three locust species. These orthologous DR-DEGs, identified independently within each species and compared across species using orthogroups, may represent phylogenetically conserved genes involved in phenotypic plasticity. Functional annotation of these shared DR-DEGs revealed genes associated with juvenile hormone signaling, cuticular proteins, and hexamerins (Supplementary Data S8). Intriguingly, several of these genes have also been implicated in caste differentiation in eusocial insects (*58*, *59*) and in horn polyphenism in beetles (*60*), suggesting that they may constitute broadly conserved “toolkit” genes repeatedly recruited during the evolution of phenotypic plasticity across insects.

To functionally assess the role of these candidate toolkit genes, we selected orthologous genes from the shared DR-DEG set that were (i) highly expressed in crowded locusts, or (ii) showed signatures of positive selection, or (iii) were supported by previous studies (*23*) (Supplementary Data S8). We designed double-stranded RNA (dsRNA) targeting these genes and conducted RNA interference (RNAi) experiments to knock down their expression in gregarious nymphs of *S. gregaria*, the species whose phase polyphenism has been best characterized (Fig. 6A). Following dsRNA design and efficiency validation, we retained five genes with robust knockdown efficiency (Supplementary Figure S12): two hexamerins-like (*hex1*, *hex2*), one inositol oxygenase-like (*miox*), one juvenile hormone acid O-methyltransferase-like (*jhamt*), and miniature gene (*unch/m*). We reasoned that these genes contribute to maintaining the gregarious phase and that their knockdown might induce a shift toward solitarious-like behavior. However, despite efficient transcript-level knockdown, none of the RNAi treatments produced significant behavioral changes (Fig. 6B).

**Figure 6:**
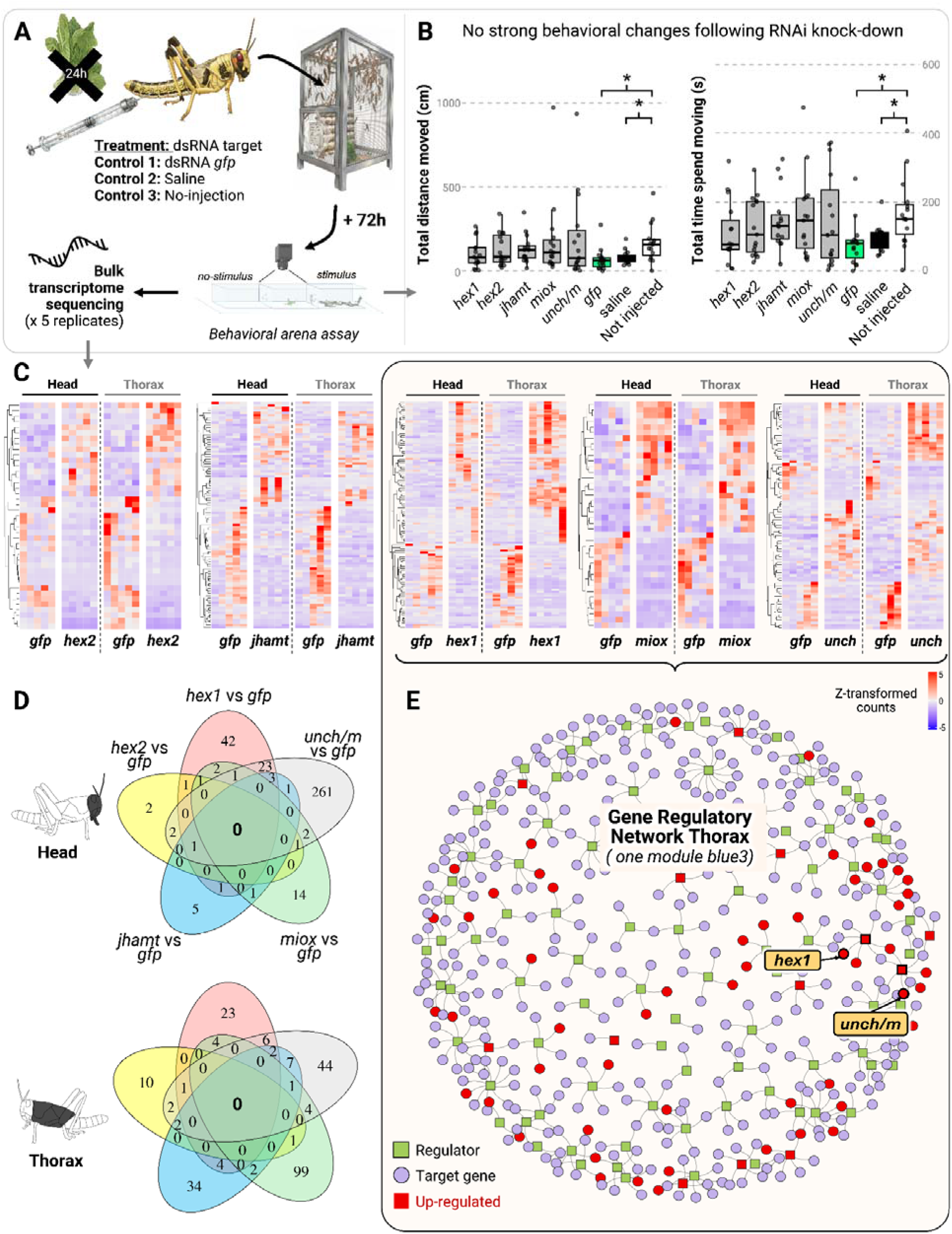
Search for candidate genes for swarming revealed the complexity of regulatory networks across tissues. Candidate genes were selected because they either a) were DEG shared by at least two locusts, b) presented strong selection pressures or c) were predicted to be key genes from the literature. Among the more extensive list, five (*hex1*, *hex2*, *miox*, *jhamt*, and *unch/m*) were successfully used for RNAi knockdown with an efficiency of >80%. Panel A presents a brief outline of RNAi design used on *S. gregaria* female gregarious nymphs to test for behavioral effects. Panel B presents some of the resulting behavioral traits measured following 72h RNAi treatment. Significant effects were only observed between controls injected vs. non-injected, rather than the direct effect of the RNAi disruption. Panel C shows that co-expression disturbance and individual variability were visible in normalized read count heatmaps. In the heatmaps, each column represents an individual used in an RNAi experiment, with high read counts in red and low read counts in blue. Panel D shows that there is no complete overlap in DEG responses in nymphs following RNAi knockdown of each gene of interest, with *gfp* as a control. Panel E displays the gene regulatory network inferred from the shared module in the thorax. This WGCNA co-expression module included *hex1*, *unch/m*, and *miox* and highlighted the upregulation of independent and multiple regulators to these genes of interest.

We then assessed whether knockdown of these candidate genes altered broader transcriptional responses by generating RNA-seq data from head and thorax tissues following dsRNA administration and comparing each treatment to a *gfp* control (Fig. 6C). We found that each knockdown induced substantial changes in gene expression, with numerous genes up- or downregulated relative to the control. In other words, these RNAi-induced transcriptional responses did not change the behavior of gregarious nymphs into solitarious ones, but they profoundly affected transcriptomic reaction norms. The expression changes associated with the RNAi treatment, while tissue-specific for each gene, were most pronounced in the thorax for *hex1* and *unch/m*. At the same time, we noted substantial individual-level variations in DR-DEGs, even among individuals reared under identical conditions and originating from the same colony (Fig. 6C). Although gene expression changes were widespread across treatments compared to the *gfp* control, we found no complete overlap among RNAi responses (Fig. 6D), indicating that knockdowns produced largely specific transcriptional perturbations. Together, these patterns suggest that a high level of intraspecific genetic variation exists within a colony and that these target genes may be downstream effector genes rather than causal drivers of locust phase polyphenism.

### Regulatory network analysis points to complex cascades of genes involved in locust phase polyphenism

To determine whether the identified DR-DEGs associated with locust phase polyphenism clustered within specific regulatory networks, we used Pfam domain predictions to annotate and infer potential regulators from genes sharing co-expression profiles in *S. gregaria*, as this is the best-annotated genome. This approach allowed us to distinguish transcription-factor-like regulators from downstream effector genes and map directional interactions. This analysis showed that each co-expressed gene was influenced by distinct upstream predicted regulators. For example, within *S. gregaria*, three candidate genes (*hex1*, *miox* and *unch/m*) co-occurred in the same thoracic module and were more highly expressed in crowded than isolated nymphs (Fig. 6E). However, the Pfam-annotated regulatory network inference showed that each of these genes was influenced by different upstream predicted regulators, indicating parallel control signals rather than a unified regulatory cascade.

Among the most compelling upstream regulators in the inferred network were the class A basic helix-loop-helix and the ecdysone-induced proteins, both of which were upregulated under crowded conditions and were predicted to directly regulate broader sets of DR-DEGs. Crucially, these candidate regulators emerged only after integrating DR-DEGs’ co-expression patterns with Pfam-derived functional annotations, highlighting the importance of combining network structure with domain-based inference. Their position upstream of *hex1*, *miox*, and *unch/m* in the inferred regulatory network suggests that key modulators of density-dependent responses may lie in early regulatory layers rather than among downstream effector genes. Rather than indicating a single hierarchical cascade, these results support a model in which multiple regulatory pathways converge on shared downstream targets. Consistent with the RNAi results, this supports the idea that our candidate genes tested here likely contribute to late-stage stabilization and maintenance of the gregarious phase rather than initiating behavioral shifts.

## DISCUSSION

Transitions from solitary living to coordinated collective behavior have evolved repeatedly across animals, reshaping how organisms avoid predators, exploit patchy resources, migrate across landscapes, and synchronize reproduction. For example, fish shoals reduce predation risk through dilution and confusion effects (*61*); eusocial insects coordinate foraging and defense through division of labor (*62*); and migratory butterflies or birds synchronize long-distance movements to track seasonal resources (*63*, *64*). In each case, collective behavior emerges from interactions among individuals who retain the capacity to behave independently, although this balance shifts toward obligate integration in eusocial systems. Swarming locusts represent one of the most dramatic examples of this shift, as density-dependent cues rapidly transform shy sedentary grasshoppers into cohesive, mobile migratory populations.

Over the past century, extensive mechanistic studies in the Migratory locust, *L. migratoria*, and the Desert locust, *S. gregaria*, have revealed key hormonal and pheromonal pathways (*9*, *65*, *66*), neuromodulatory pathways (*67*), and epigenetic pathways (*34*, *68*, *69*) involved in phase transition. However, most of these investigations have focused on single-species, leaving unresolved whether the described mechanisms represent conserved molecular blueprints for locust swarming or lineage-specific solutions. Given that locust species have independently evolved and differ in ecological context, outbreak dynamics, and density thresholds, their plasticity and collective behavior may arise through distinct evolutionary routes. By integrating chromosome-scale genomics, comparative transcriptomics, behavioral assays, and functional perturbation across six *Schistocerca* species, our study shows that similar swarming phenotypes have emerged from largely distinct molecular architectures. At the level of genome structure, we detected no singular “locust signature”. Chromosome organization, gene content, and large-scale synteny remain broadly conserved across the genus. Gene family expansions and signatures of positive selection reflect lineage-specific evolutionary histories rather than a shared genomic innovation uniquely associated with swarming. Thus, similarities in collective behavior do not appear to originate from a structural genomic transformation.

Instead, divergence is apparent at the level of regulatory responsiveness. We identified a spectrum of density-dependent behavioral plasticity across *Schistocerca*, paralleled by a spectrum of density-responsive transcriptional remodeling. Species exhibiting strong density-dependent behavioral shifts also show extensive differential gene expression, whereas species with reduced behavioral plasticity display limited transcriptional responsiveness. What distinguishes locusts from closely related grasshoppers is therefore not genome architecture, but the scale and organization of density-responsive regulatory programs. Strikingly, however, we also discovered that different locust species, even within the same genus, have evolved different density-responsive genes to achieve a similar expression of locust phase polyphenism. More intriguingly, these species-specific density-responsive genes were partially involved in similar functions, suggesting that there may be functional constraints on achieving phenotypic plasticity (*70*), even though the genes themselves may differ. Despite the low overlap in density-responsive genes among locust species, a small number of their shared genes have been implicated in phenotypic plasticity of other insect systems (*59*, *60*, *71*). In other words, these genes may represent toolkit genes that are redeployed across insects that evolved phenotypic plasticity. However, our functional and regulatory network analyses indicate that these genes act as downstream effectors within distributed gene networks rather than as master regulators of behavioral plasticity.

Our biologically integrative and phylogenetic approach indicates that locust phase polyphenism reflects evolutionary convergence built on flexible regulatory architectures. While understanding gene regulatory networks has been shown to be key for uncovering the origins and evolution of phenotypic diversification in plastic traits across taxa (*72*, *73*), the large and highly interconnected networks observed in locusts present both a challenge and an opportunity for identifying the regulatory logic underlying this complex trait. An ancestral capacity for density-dependent responsiveness may have provided a shared substrate within *Schistocerca*, while subsequent lineage diversification produced species-specific transcriptional and network configurations. Collective behavior in the form of swarming thus arises not from conserved genes, but from the repeated assembly of regulatory systems that satisfy common ecological and physiological demands. More broadly, our results challenge the assumption that convergent phenotypes often result from convergent molecular mechanisms (*74*). Complex behavioral transitions may evolve through parallel reconfiguration of gene regulatory networks constrained by functional necessity but free in their specific genetic implementation. Locust swarming, therefore, provides a model for understanding how distributed genomic architectures generate emergent collective states, illuminating general principles of phenotypic plasticity and the evolution of coordinated behavior across animals.

With the reference genomes and integrated behavioral and transcriptomic datasets presented here, we move closer to identifying the proximate mechanisms that differentiate swarming locusts from their sedentary relatives. Nevertheless, important questions remain. The initiation of phase transition likely involves early epigenetic modulation, dynamic gene regulatory interactions, and cell-type specific responses that are not fully resolved in bulk genomic and transcriptomic analyses. Early-responsive genes that initiate phase transition may act transiently in specific tissues or cell types and could be masked in bulk transcriptomic analyses by compensatory dosage mechanisms that stabilize downstream expression profiles (*34*, *75*). Dissecting these processes at finer spatial and temporal scales will be essential to uncover the initial molecular triggers leading to collective behavior and to determine how distributed regulatory systems coordinate rapid phenotypic transformation. Beyond advancing evolutionary theory, understanding the genomic architecture of locust phase polyphenism has practical implications. Swarming locusts remain among the most destructive agricultural pests worldwide, and improved knowledge of the regulatory networks underlying gregarization may ultimately inform more precise and sustainable management strategies. By establishing a comparative genomic framework across swarming locust and non-swarming grasshopper lineages, this study provides a foundational context for future mechanistic and applied investigations of plasticity and collective behavior.

## MATERIALS AND METHODS

### Sampling for genome assembly

We acquired samples from six *Schistocerca* species, including locusts (*S. gregaria*, *S. cancellata,* and *S. piceifrons*) and closely related non-swarming grasshoppers (*S. americana*, *S. serialis cubense,* and *S. nitens*), which show various degrees of density-dependent phenotypic plasticity (*4*, *23*). Live female specimens of every species were collected from long-term crowded colonies maintained at lab-rearing facilities at either the Department of Entomology at Texas A&M University (USDA PPQ 526-24-280-15160), the School of Sustainability at Arizona State University, or the International Center of Insect Physiology and Ecology (ICIPE) in Kenya. Except for *S. gregaria,* which was obtained at the ICIPE after one generation, specimens were acquired after at least five generations following lab colony establishment. We sourced *S. gregaria* specimens collected from outbreak regions in Kenya during the 2020 upsurge. At Texas A&M University, the *S. piceifrons* colony originated from an outbreak population in Yucatan near Tizimin, Mexico, collected in October 2015 and imported under the USDA permit APHIS PPQ P526P-15-03851. The *S. americana* colony was established from a population in Brooksville, Florida, collected in September 2010. The *S. serialis cubense* colony originated from a population in Islamorada in the Florida Keys collected in January 2011. The *S. nitens* colony was established from a population in Terlingua, Texas, collected in May 2015. At Arizona State University: the *S. cancellata* colony originated from an outbreak population in Catamarca province, Argentina, collected in 2016. At ICIPE, the *S. gregaria* specimens (F1) originated from a mated pair collected near Samburu, Kenya, in October 2020. Two females per species were snap-frozen and stored at -80°C until laboratory processing for PacBio HiFi and Hi-C library preparation.

### DNA extraction and whole genome sequencing

To address the challenge posed by the large genome size in Orthoptera (*24*, *76*), we have chosen a strategy combining high-fidelity (HiFi) long reads generated using Pacific Biosciences (PacBio) sequencing, combined with chromatin conformation capture Hi-C data (*77*). Snap-frozen whole-body samples were exchanged between facilities: USDA-ARS in Hilo (Hawaii) for PacBio sequencing and Baylor College of Medicine in Houston (Texas) for Hi-C sequencing. Among the two adult females per species, the one yielding the better high-molecular-weight DNA was used for both HiFi and Hi-C library preparation. For each species, approximately 5 μg of DNA was used to create PacBio HiFi sequencing libraries using the SMRTbell Express Template Prep Kit 2.0, following standard low-input DNA procedures. Fragments shorter than 3 kb were removed via AMPure size selection. The resulting libraries were sequenced on a PacBio Sequel IIe system using 12 to 18 SMRT Cell 8M runs per species, achieving >27× genome coverage (Table X1). Subsequently, hind leg tissue was prepared for Hi-C, consistent with the standard protocols used by the DNA Zoo consortium, and *in situ* Hi-C was then performed following the approach of (*78*). The resulting Hi-C libraries were sequenced to ∼10× coverage using short paired-end reads on an Illumina NovaSeq 6000 (Table X1).

### Genome assembly

We used PacBio HiFi long-read data to generate the initial genome assemblies at the contig level. Prior to the assembly of each dataset, raw HiFi reads were filtered using HiFiAdapterFilt (*79*) to remove any circular consensus sequencing (CCS) reads that contained PacBio adapter sequences. The filtered data were assembled using HiFiASM (v. 0.15.4) (*80*) using default parameters in diploid mode. The assembled PacBio HiFi contigs were then placed in the correct order and orientation to form chromosome-length scaffolds using *in situ* Hi-C data and 3D-DNA for automated scaffolding (*81*), followed by Juicebox Assembly Tools for assembly finishing (*82*). Chromosome assignments were done for each species using previously assessed karyotypes for *Schistocerca* as a guide (*36*). Hi-C contact maps for each species are available using an interactive interface hosted on DNA Zoo (e.g., for *S. piceifrons* https://www.dnazoo.org/assemblies/schistocerca_piceifrons).

### Density-dependent transcriptome sequencing

To identify and compare density-dependent responsive genes, we sequenced the transcriptomes of long-term crowded and one-generation isolated individual specimens of the six *Schistocerca* species, as described by Foquet et al. (*23*). A total of 80 transcriptomes had been previously generated for the locust *S. piceifrons* and the grasshoppers *S. americana*, *S. nitens,* and *S. serialis cubense*, accessible through the NCBI BioProject PRJNA633949. These transcriptomes were independently generated from whole heads or thoraxes. We produced comparable RNA-sequencing datasets for last-instar female nymphs (around 72h after molting) from the two remaining locusts, *S. gregaria* and *S. cancellata,* reared using the methods described by Foquet et al. (*23*) and Pocco et al. (*83*).

#### Rearing regime

The *S. gregaria* colony originated from a long-term inbred and well-characterized laboratory population maintained at the University of Leicester, initially obtained from a local supplier in the United Kingdom, and subsequently transferred to Arizona State University in 2020. Later, in 2021, the colony was imported and established at Texas A&M University under the permit USDA APHIS PPQ P526P-19-02151. The *S. cancellata* colony originated from an outbreak population in Catamarca province, Argentina, collected in 2016 and was maintained at the Centro de Estudios Parasitologicos y de Vectores (CEPAVE) in La Plata, Argentina (*83*). The behavioral assays and RNA-sequencing sampling were conducted at Texas A&M University for *S. gregaria* and at the CEPAVE for *S. cancellata* at least two generations after establishment. For both species, we reared the insects under two density conditions following standard methods developed for *Schistocerca* (*29*): 1) a crowded condition to induce gregarious phenotypes through exposure to high conspecific density and 2) an isolated condition to induce solitarious phenotypes, in which a single individual developed without sensory contact with conspecifics (*22*, *31*). To induce gregarization, approximately 100 nymphs of *S. cancellata* were reared in a wire-screened, ventilated aluminum cage (30 x 22 x 22 cm) and fed daily fresh lettuce, wheat bran, and cabbage. For *S. gregaria*, about 300 nymphs were reared in slightly larger cages (30.5 x 30.5 x 30.5 cm) and fed daily with fresh lettuce, wheat grass, and wheat bran. To induce solitarization, hatchlings originating from the long-term crowded stock were individually reared in custom-designed cages (10.16 x 10.16 x 25.4 cm) until reaching the final nymphal instar. These individual cages, located in a separate room from the crowded colonies, were visually shielded and provided with positive, filtered airflow from an air compressor to prevent visual, tactile, and olfactory stimuli from conspecifics. Isolated nymphs were fed *ad libitum*, following the same dietary regime as crowded nymphs, with food replaced every 1-3 days. In both density conditions, insects were reared under controlled photoperiods: 12h of light and 12h of darkness at Texas A&M University (30°C) and 14 h of light and 10 h of darkness at the CEPAVE (30°C).

We targeted female nymphs approximately 72h after molting to their last nymphal instar as a standard approach (*23*, *83*). While tracking the molting status of isolated nymphs is straightforward with daily and frequent observation, it is challenging for densely crowded cages with nymph cohorts molting at different times of day. We addressed this issue by marking newly molted last instar nymphs in isolated and crowded conditions using water-acrylic-based pens (*Posca*) on the thorax, using a daily cut-off time at noon, and tracking them every 4-5 hours. Similar to observations made by Foquet et al. (*23*), we did not note a change in behavior or mortality rates following body marking. We randomly sampled 5 to 6 last instar nymph females per condition and performed live dissections with sterilized tools in the morning between 9:00 am and 12:00 pm. Upon dissection (< 3 mins), head (including antennae) and thorax tissues (including wings but not legs) were snap-frozen for *S. gregari*a or preserved in RNAlater (*Qiagen*) for *S. cancellata* and kept at -80°C until RNA extraction.

#### RNA-sequencing

The nucleic acid extraction, quantification, library preparation, and sequencing for *S. cancellata* followed the same workflow as the one previously described for the four species *S. americana*, *S. piceifrons*, *S. nitens,* and *S. serialis cubense*, for which RNAseq datasets were previously generated by Foquet et al. (*23*). Minor changes were made for *S. gregaria* to reflect technological improvements, as its tissue sampling was completed last. In brief, each of the 24 tissues (six biological replicates per condition) was individually ground using a Tissue Lyzer II (*Qiagen*). We extracted the total RNA with the simplyRNA Tissue kit and the automated Maxwell RSC instrument (*Promega*) following the manufacturer’s recommendations. RNA concentration was quantified using a Qubit 4.0 fluorometer with the RNA Assay Broad Range Kit (*Invitrogen*), while RNA quality was assessed using TapeStation RNA Screen Tape (*Agilent*). Whole transcriptome library preparation included a ribosomal RNA depletion step using the Illumina Stranded Total RNA with Ribo-Zero kit (*Illumina*) following the manufacturer’s recommendations. Equimolar pooled, paired-end libraries were sequenced on a single NovaSeq SP PE150 flow cell at the TIGGS Molecular Genomics Core at Texas A&M University. Demultiplexing and initial quality control were performed and provided by the sequencing center. We note that ribosomal RNA depletion using the commercial Ribo-Zero kit was suboptimal for *Schistocerca* species, resulting in residual rRNA contamination. This was mitigated by increased sequencing depth (Supplementary Data S9).

### Isoform sequencing of *S. gregaria*

To sequence full-length isoforms, RNA extraction, PacBio IsoSeq library prep, and sequencing was performed from dissected brain tissue, testes, fat bodies, midgut, mouth parts, and antennae from a single male, and ovaries from a single female. Ribonucleic acid extraction was performed using Trizol and the Zymo Direct-zol 96 MagBead RNA kit. cDNA was synthesized from the RNA using the NEBNext Single Cell/Low Input cDNA Synthesis and Amplification kit using the PacBio Iso-Seq Express Template Switching Oligo. The Iso-Seq library was prepared from cDNA with the PacBio Iso-Seq Express Library Preparation Using SMRTbell Express Template Prep Kit 2.0. Sequencing was performed on a Sequel II System using Binding Kit v2.0, Sequencing kit v2.0, and SMRT Cell 8M. To target HiFi reads, three libraries were barcoded individually and pooled for sequencing using a 30 hour movie time on one SMRTcell. Raw subreads were converted to HiFi data by processing with CCS to call a single high quality consensus sequence for each molecule, using a 99.5% consensus accuracy cutoff.

### Genome annotation and quality assessment

We assessed the quality of the draft assemblies at each step by measuring the total length of the assembly, the number of contigs and their lengths, and the contigs and scaffolds’ N50 values using AGAT and Gfastats (*84*, *85*) (Supplementary Table S1). Each polished assembly with chromosome assignments was then uploaded to the NCBI genome database, filtered, and cleaned of minor microbial contaminants. All six draft genomes were then annotated using the NCBI Eukaryotic Genome Annotation Pipeline (EGAP) (*86*) and were publicly released as reference genomes (RefSeq) under the umbrella BioProject PRJNA772266. Additionally, the six genomes are available on the genome browser of the USDA i5k Workspace, enabling manual genome annotation. Details on the origin of each RNA-seq dataset used to support genome annotation are available in the NCBI SRA archive and the annotation reports for each species. The annotation of *S. gregaria* benefited from the largest transcriptomic evidence using EGAP with both ∼1.70 billion RNA-seq short reads (using STAR (*87*)) and ∼4.91 million IsoSeq reads (using minimap2 (*88*)). Annotation was performed after repeat masking, in which repeat elements were first detected using RepeatModeler (*89*) and then masked using RepeatMasker (*90*). The completeness of each genome was estimated using 3,114 core insect genes from the insecta_odb12 database with BUSCO v.4.1.4 (*91*), to produce final gene and feature statistics. We used BlobToolKit (*92*) and OMark (*93*) to visualize and summarize genome assembly statistics regarding genome coverage, scaffolds/chromosomes length, and gene repertoire completeness.

### Orthologs detection

We identified orthogroups (i.e., gene families) and orthologs of single-copy genes by comparing the six *Schistocerca* genomes with Orthoptera (n = 4) and Polyneoptera (n = 3) outgroup genomes using OrthoFinder (v2.5.5) (*49*). Outgroups were selected based on phylogenetic proximity, large genome sizes, the use of hybrid assembly techniques, the availability of genome assemblies on NCBI (preferably at the chromosome level), and the presence of corresponding annotation files (last NCBI search: 28 April 2024). The resulting orthopteran outgroup species were the migratory locust *Locusta migratoria* (GCA_026315105.1), the Mormon cricket *Anabrus simplex* (GCF_040414725.1), the long-cercus field cricket *Gryllus longicercus* (GCA_038098605.1), and the two-spotted field cricket *G. bimaculatus* (GCA_017312745.1). The resulting polyneopteran outgroups included *Dryococelus australis* (GCA_029891345.1), the European stick insect *Bacillus rossius redtenbacheri* (GCF_032445375.1), the drywood termite *Cryptotermes secundus* (GCF_002891405.2), and the American cockroach *Periplaneta americana* (GCF_040183065.1). For the *Gryllus* outgroups, RefSeq annotations were not available, and we used and formatted user annotation files using AGAT (*84*). For *L. migratoria*, we annotated the genome using the EGAPx pipeline and based on 13 RNA-seq datasets from NCBI SRAs generated from gregarious and solitarious nymphs. We followed the workflow by Bardkull and Moreau (*94*) to select the longest isoform of each gene using the R “orthologr” package, before cleaning and preparing the data for direct homologous gene analysis. In OrthoFinder, we conducted a sequence search with DIAMOND, using IQTREE for gene and species tree inference, MAFFT for multi-species alignment, and an MCL inflation parameter of 2 without specifying a species tree prior. The species tree was generated using the msa option, which used a concatenated multiple sequence alignment from solely single-copy orthologs (*95*). Orthogroups and single-copy genes were extracted based on the OrthoFinder results and were subsequently used for differential gene expression, gene family contraction and expansion, and signature of selection analysis.

### Repeat landscape analysis

The repeat content files were generated by first running all six *Schistocerca* genomes through RepeatModeler (Flynn et al., 2020) and then RepeatMasker (Smit et al., 2013) to identify and denote repetitive sequences within the genomes. We then used the calcDivergenceFromAlign.pl script from the RepeatMasker module to generate a distance file which contains the annotated base counts from each alignment. Using a version of the distance file containing only the Kimura scores, we then generated the repeat landscape figures in R using a script named kimura-plot (*96*).

### Synteny analysis

We first ran our six *Schistocerca* genomes and Polyneoptera outgroup genomes through the FormicidaeMolecularEvolution workflow (*94*) to select the single longest gene isoform in each genome, accounting for gene duplication and decreasing the computational time of OrthoFinder. These refined files were then used as input for OrthoFinder to detect orthologous genes and generate alignment files for each genome against the 11 other species included in the run. We then used GENESPACE (*97*) to generate a synteny plot for all six *Schistocerca* genomes with *S. gregaria* as the reference genome to characterize the structural similarity of the orthologous genes identified by OrthoFinder.

### Comparative genomics analysis

#### Gene family evolution

We used the gene family analyzer CAFE5 (*50*) to explore whether the evolutionary history of swarming locusts differed from that of sedentary grasshoppers regarding gene gain and losses. We tested all six genomes of *Schistocerca* and the seven Polyneoptera outgroups by formatting the total gene count matrix per orthogroups output generated by OrthoFinder with custom scripts. To avoid model convergence failures and to ensure accurate estimation of the global birth-death (λ) across each gene tree, we filtered gene families presenting extreme copy number variation. Thus, we removed orthogroups with more than 400 genes in total or a size variation greater than 100 across species. Then, we transformed the best-rooted species tree inferred by Orthofinder and IQTREE into an ultrametric tree using ‘make_ultrametric.py.’ The parameter λ was estimated without assuming gene family rate variation, and the results were visualized using CafePlotter.

#### Episodic positive selection analysis

To assess how selective regimes have shaped gene evolution in *Schistocerca* and to investigate species-specific adaptations in swarming lineages, we used three different modules from the Hyphy v2.5.71 package (*98*) to compute and test the dN/dS ratio. For phylogenetic reconstruction, we used 3,128 concatenated nuclear gene protein sequences from single-copy orthogroups. We recovered the monophyletic relationship of *Schistocerca* with only a difference of branch placement between *S. nitens* and *S. cancellata* compared to the previous time-calibrated tree (*4*). Selection tests were performed using aBSREL (adaptive Branch-Site Random Effects Likelihood) (*54*), BUSTED (Branch-Site Unrestricted Statistical Test for Episodic Diversification) (*98*), and RELAX methods for relaxed selection (*55*) on single-copy orthogroup sequence alignments and gene trees to detect signals of positive or lineage-specific selection. For both aBSREL and BUSTED, we conducted analysis under two configurations: 1) unlabeled phylogenies for exploratory analysis, and 2) labeled trees in which branches corresponding to the phenotype of swarming locust species (*S. gregaria, S. piceifrons, S. cancellata,* and *L. migratoria*) were designated as “foreground” to determine whether lineages experienced heightened selective pressures relative to other clades. RELAX requires a labeled phylogeny. We used aBSREL to identify if specific branches experienced episodic diversifying selection in each gene family. We used BUSTED to test for alignment-wide episodic selection across the 13 genomes by asking whether a gene has experienced positive selection for at least one site on at least one branch (*99*). We used RELAX to test for relaxed or intensified selection on all orthogroups that were found to be under positive selection from aBSREL and BUSTED runs. RELAX is not designed to detect positive selection but rather to determine the selection intensity parameter (k), where values of k > 1 indicate intensified selection and k < 1 indicate relaxed selection in foreground lineages. In all cases, we tested each multispecies orthogroup defined by OrthoFinder using default parameters and allowing synonymous rate variation with the flag ‘–srv Yes’. The results were compiled from all JavaScript Object Annotation (JSON) file outputs using a slightly modified R script from Bardkull and Moreau (*94*) to perform Benjamini-Hochberg (BH) false discovery rate (0.05) corrections on the p-values reported.

### Demographic inferences with PSMC

We inferred long-term demographic histories for each species using the Pairwise Sequentially Markovian Coalescent model (PSMC), which reconstructs changes in effective population size (Ne) through time from a single diploid genome (*100*). We used PSMC to explore whether species sharing swarming and outbreaking traits would present distinct and similar changes in population sizes from sedentary grasshoppers in response to long-term habitat changes. We followed the protocols described by the PSMC package developer’s GitHub (https://github.com/lh3/psmc). For each species, we mapped the paired-end Hi-C reads back to the corresponding reference genome using the BWA aligner. Alignments were converted to BAM format, coordinate-sorted, and used to generate a diploid consensus sequence using bcftools. We combined ‘bcftools mpileup’ and ‘call’ commands, followed by the executable script ‘vcftutils.pl’ with default parameters except for minimum base and mapping quality thresholds set at 20. We then converted the consensus fastq files into PSMC input format using ‘fq2psmcfa’ and ran PSMC with the following parameters: -N25 -t15 -r5 -p “4+25*2+4+6”. For each species, we generated 100 bootstrap replicates after splitting the input with ‘splitf’.

Since no Orthopteran and more specifically Acrididae genome-wide mutation rate has been reported so far, we scaled the PSMC trajectories for each *Schistocerca* species by using the mutation rate estimated for the model insect *Drosophila melanogaster* (2.8 x 10 mutations per site per generation) (*101*). Because generation time varies among locust and grasshopper species and can change with ecological conditions, we explored a range of generation-time scenarios from one to three generations per year for all species (*18*, *102*). For the main comparative plots, we used two generations per year (g = 0.5). We visualized the bootstraps trajectories and multi-species comparisons using ‘psmc_plot.pl’ and custom R scripts.

### Comparative transcriptomics analysis

#### Read mapping and gene count

We retrieved paired-end fastq files for *S. piceifrons*, *S. americana*, *S. nitens,* and *S. serialis cubense* from NCBI BioProject PRJNA633949, using SRA-Toolkit v2.10.9 and the fasterq-dump --split-3 parameter. We developed a Snakemake pipeline to automate all downstream analysis. Both downloaded and newly generated reads were assessed for quality using FastQC (*103*), and summary reports were compiled using MultiQC (*104*). We then subjected all transcript reads to quality filtering and adapter trimming using fastp v0.23.2 (*105*), retaining reads with a minimum length of 50 bp. To perform cross-species comparisons and predict gene expression profiles associated with conspecific density changes, we used two alternative mapping approaches, as translating transcriptomics results across species can be challenging. In the first method, referred to as the *“shared RefSeq*” strategy, we leveraged the close phylogenetic relationships among *Schistocerca* species, with *S. gregaria* occupying a position closest to the last common ancestor of the genus, consistent with its biogeographic distribution and origin (*4*, *106*). We, therefore, mapped all reads to the annotated *S. gregaria* genome, allowing direct comparison of gene identifiers across species. However, this strategy has the drawback of potentially falsely reflecting true transcript abundance and missing lineage-specific or rare transcripts due to sequence divergence or absence in the reference genome (*107*, *108*). In the second method, referred to as the “*species-specific RefSeq*” strategy, we mapped transcript reads from each species to its own annotated RefSeq genome. Mapping to individualized transcriptomes has been shown to increase read-mapping accuracy and improve transcript abundance quantification (*107*), but this method complicates direct gene identifier comparisons across species. Cross-species transcriptome studies can address this problem by focusing on core sets of genes that are 1:1 orthologs (*109–111*). As suggested by the OrthoFinder outputs, *Schistocerca* genomes showed many large-scale duplication events. To account for this, we performed our analyses at the orthogroup level, including not only 1:1 orthologs but also orthogroups containing 1:N and N:1 relationships.

#### Differentially expressed genes

For both “*shared RefSeq*” and “*species-specific RefSeq*” strategies, we indexed each genome with the STAR aligner v2.7.10b (*87*, *112*) with the parameters ‘--sjdbOverhang 149’ and ‘--alignIntronMax 2500000’, the latter based on manual inspection of gene annotations. We then aligned reads to the reference genome using the 2-passes mode of STAR and by outputting raw gene counts with the options ‘--quantMode TranscriptomeSAM GeneCounts’. Following exploratory analysis, we opted for featureCounts from Subread v2.0.3 (*113*) to extract per-sample gene and transcript counts. These files were then used as input for differential expression gene (DEG) analysis with the R package DESeq2 (*114*) using the function DESeqDataSetFromHTSeqCount. We conducted a preliminary bioinformatic ribosomal RNA depletion in the gene count matrices by excluding “rRNA” features from the .gff file. The workflow was run independently for the thorax and head tissues and each species. Upon data exploration using PCA and a sample distance matrix, outliers were removed before computing final DR-DEGs using DESeq2’s variance stabilizing transformation. For downstream analysis, only DR-DEGs with a minimum absolute log_2_-fold change > 1 and corrected p-value (BH corrected) < 0.05 were kept. Visualization of DEG results through heatmaps, volcano plots, and MA plots was all done in R.

#### Selection of candidate genes

Summary and overlap analysis for cross-species transcriptomic results resulting from both mapping strategies was conducted in R with the VennDiagram and UpSetR packages. The only difference for the “*species-specific RefSeq*” approach is that DEG identifiers were linked to orthogroup identifiers from the OrthoFinder output. We extracted overlapping DEG identifiers to find genes shared among: 1) the three swarming locusts (*S. gregaria, S. piceifrons*, and *S. cancellata)*, 2) the locusts and the intermediate phenotypically plastic species *S. americana,* and 3) all *Schistocerca* species. The aim was to identify a possible core set of genes associated with swarming behavior and its underlying regulatory mechanisms.

### Gene co-expression clustering and regulatory network inferences

#### Weighted gene co-expression network analysis

To determine whether density-responsive genes formed coordinated transcriptional modules associated with behavioral traits, we performed weighted gene co-expression network analysis (WGCNA) separately for each locust species and tissue (head and thorax). We used variance-stabilized expression matrices derived from DESeq2 and paired them with behavioral trait tables containing the median values for each species and rearing condition for the main behavioral variables measured in the arena assays, including moved distance, time spent in the stimulus zone, time spent in the non-stimulus zone, and time spent on the stimulus wall. Because behavioral measurements were obtained at the group level for each condition, median trait values were used to represent each experimental group, allowing integration with transcriptomic profiles while minimizing the influence of outliers. For *S. gregaria*, which had substantially more behavioral trials than the other species, we randomly subsampled 50 isolated and 50 crowded trials to make the behavioral input comparable across species.

Before network construction, we verified that expression matrices and trait tables were aligned by sample identity and converted non-numeric trait variables into numeric format when required. We selected a soft-thresholding power using the scale-free topology criterion implemented in pickSoftThreshold and used a power of 12 for the final networks. We then identified co-expression modules with blockwiseModules in WGCNA (*115*) using an unsigned network, a minimum module size of 30 genes, no reassignment threshold, a merge cut height of 0.2, and a maximum block size of 20,000 genes. We calculated module eigengenes and tested their correlations with behavioral traits using Pearson correlations and Student’s asymptotic p-values. We summarized module size, module–trait relationships, and eigengene structure with heatmaps and network visualizations generated in R (*116*). Hub genes were identified within focal modules using module membership scores.

#### Transcription factor prediction

For each *Schistocerca* species, we mapped the predicted protein sequenced from NCBI against the Pfam database (*117*) and HMMER (*118*) to identify proteins containing conserved domains associated with transcription factor families. Using a custom script, we filtered the HMMER output to retain a curated set of transcription factor-associated Pfam domains and reformatted the results to include the Pfam ID, NCBI protein accession number, and domain annotation.

For *S. gregaria*, we used these predicted regulators together with the expression matrix of the focal tissue to infer gene regulatory networks in R with the BioNERO package (*119*). Specifically, we constructed a SummarizedExperiment object from the normalized expression data, provided the filtered list of predicted regulators as input to ‘exp2grn,’ and identified hub regulators with ‘get_hubs_grn.’ Hub genes were identified based on network connectivity. We visualized the resulting regulatory networks with BioNERO and used these networks to distinguish upstream transcription factor-like candidates from downstream effector genes within co-expression modules associated with locust phase polyphenism.

### Functional validation with RNAi

#### Double stranded RNA synthesis and efficiency testing

For dsRNA synthesis, total RNA was extracted using the method described in the RNA-seq section above. The extracted RNA was reverse transcribed into cDNA using the iScript cDNA synthesis kit (Bio-Rad) according to the manufacturer’s instructions. Gene-specific fragments were amplified by PCR using primers (Supplementary Table S2). PCR amplification was performed under the following thermal cycling conditions: 94 °C for 30 s, 55–60 °C for 30 s and 72 °C for 1:30 min for 10 cycles. This was followed by 25 cycles 94 °C for 30 s, 69 °C for 30 s, and 72 °C for 1:30 min, with a final extension at 72 °C for 2 min. PCR products were purified using the Monarch PCR and DNA Clean Up Kit (New England Biolabs) according to the manufacturer’s protocol. The purified amplicons were used for dsRNA synthesis using the MEGAscript RNAi Kit (Thermo Fisher Scientific) following the manufacturer’s instructions and the previous protocol established for *Schistocerca* (*120*). For non-target control in experiments, dsRNA of *gfp* was synthesized using a TOPO GFP plasmid (Addgene). To determine RNAi efficiency, freshly molted last instar nymphs were injected with 1 μg of dsRNA targeting genes (*hex1, hex2*, *jhamt*, *miox*, *unch*/*m*), while dsRNA targeting *gfp* was used as a non-target control. The nymphs were starved for 24 h before the experiments. For each target gene, both male and female nymphs were included to account for potential sex-specific differences. Injections were done into the 2nd abdominal segment using a Hamilton microsyringe (700 series, 705RN, 50 µl, Sigma Aldrich) with a 22-gauge needle. Seventy two hours after injection, head and thorax tissues were dissected, snap-frozen in liquid nitrogen, and stored at -80 °C for subsequent RT-qPCR and RNA-seq analysis. qPCR amplification was performed on a CFX Connect Real Time PCR System (Bio-Rad) using gene-specific primers (Supplementary Table S2). Each qPCR reaction was performed with two technical replicates and three to ten biological replicates and contained 10 µl of SYBR Green supermix (Bio-Rad), 500 nM primers, and nuclease free water to make up the volume of 20 µl. The thermal cycling included initial denaturation for 3 min at 95 °C, followed by 40 cycles at 95 °C for 10 s and 60 °C for 30 s. No template control was included to check for reaction mixture contamination. Relative mRNA levels were analyzed using CFX Manager 2.1 (Bio-Rad) with the 2^-ΔΔCT^ method. The results were normalized using two reference genes, elongation factor 1 alpha (*ef1*) and ubiquitin-conjugating enzyme 10 (*ubc10*). For RNA-seq, total RNA was extracted from the head and thorax tissues using the same method described in the RNA-seq section above. A total of 60 RNA libraries were prepared from whole head and thorax tissues (5 biological replicates per tissue per gene) using the Illumina stranded Total RNA with Ribo-Zero kit (Illumina) according to the manufacturer’s instructions. Equimolar pooled libraries were sequenced on a single lane of the NovaSeq XPlus PE 150 at Novogene (Sacramento, CA).

#### RNA interference treatments

The *S. gregaria* colony was maintained at Texas A&M University under controlled laboratory conditions using standard methods (*29*). RNA treatment followed the same workflow as for the RNA efficiency determination protocol described above. Target genes included hexamerins (*hex1*, *hex2*), juvenile hormone methyltransferase (*jhamt*), inositol oxygenase (*miox*), and an uncharacterized gene later annotated as *miniature* (*unch/m*). Control treatments consisted of locust saline injections, ds*gfp* injections, and non-injected individuals. Injection needles were sterilized immediately prior to use by sequential immersion in acetone, 100% ethanol, and locust saline. After injection, gregarious nymphs were returned to their original rearing conditions and maintained under standard environmental parameters. For each injected gene, five female specimens were snap-frozen following behavioral trials and subsequently stored at -80°C. Behavioral assays were conducted 72 h post-injection, corresponding to four days post-ecdysis.

#### Behavioral assays

Behavior phase state was quantified using a rectangular Roessingh-style arena (57 × 31 × 11cm) designed to assess locomotor activity and attraction to conspecifics (*32*). The arena consisted of a central test area flanked by two end chambers separated by perforated barriers that permitted visual and chemical cues but prevented physical contact. One end chamber contained a stimulus group of 50 crowd-reared final instar nymphs, while the opposite chamber remained empty to serve as a non-stimulus side. Test locusts were introduced into the arena via a darkened 50 mL syringe and held for approximately 2 min prior to release to minimize handling stress and standardize trial initiation. Sex of focal individuals and stimulus sides were balanced within treatments. Trials were recorded from above using a high-resolution Basler camera (acA1300-60gc, 1280x1024 pixels, 30 frame/s). The first 10 minutes of behavior were tracked and analyzed using EthoVision XT 17 software (Noldus Information Technology Inc., Leesburg, VA, USA). Locomotor tracks were smoothed using a 0.2 cm minimum-distance-moved filter to reduce tracking noise. After each trial, the arena was cleaned with ethanol to remove residual chemical cues. The arena floor was divided longitudinally into three equal zones: a stimulus zone adjacent to the chamber containing conspecifics, a non-stimulus zone at the opposite end of the arena, and a central middle zone. Each zone included the corresponding arena walls. EthoVision XT was used to extract the following variables: total distance moved (cm), total time spent moving (s), and time spent in each zone (s). These measures are widely used to characterize locust phase state, with increased locomotor activity and attraction to conspecifics indicative of gregarious behavior, and reduced movement and avoidance of the stimulus zone characteristic of solitarious behavior, consistent with previous studies of locust phase polyphenism. Statistical analyses were performed with R (*116*). For each behavioral variable, normality was assessed within treatments using Shapiro-Wilk tests. Treatments were then compared to saline-injected controls (n = 15 per treatment) using two-tailed Welch’s tests when assumptions of normality were met, and Wilcoxon rank-sum tests for non-normally distributed data.

#### DEG analysis following RNAi knockdown

Differential gene expression analyses were performed following the same read processing, mapping, and quantification pipeline described in the comparative transcriptomics section above. For RNAi experiments, we compared each gene knockdown condition (*hex1*, *hex2*, *jhamt*, *miox*, *unch/m*) against the *gfp* control using a design formula accounting for treatment effects. Analyses were conducted separately for each tissue (head and thorax). Genes were considered differentially expressed if they showed an adjusted p-value (Benjamini-Hochberg correction) < 0.05 and an absolute log_2_ fold change > 1. To identify shared and treatment-specific transcriptional responses, we compared sets of differentially expressed genes across RNAi conditions and tissues. Overlapping genes were identified using set-based comparisons, and results were visualized using intersection plots and heatmaps generated in R (*116*).

### Functional annotation and enrichment analyses

To functionally characterize genes identified in differential expression, gene family expansion, and positive selection analyses, we performed GO and KEGG enrichment analyses using annotations derived from EggNOG-mapper (*121*). Protein sequences were annotated against the EggNOG database using default parameters, and corresponding GO terms and KEGG pathway assignments were retrieved for downstream analyses.

For each gene set of interest (e.g., differentially expressed genes, expanded/contracted gene families, or genes under positive selection), enrichment analyses were conducted by comparing the frequency of functional categories in the focal gene set against a background set consisting of all annotated genes in the corresponding species. Statistical significance of enrichment was assessed using Fisher’s exact test, and resulting p-values were corrected for multiple testing using the Benjamini–Hochberg false discovery rate (FDR) procedure. Terms with adjusted p-values below 0.05 were considered significantly enriched.

To reduce redundancy and facilitate interpretation of GO results, enriched GO terms were summarized using REVIGO (*122*), which clusters semantically similar terms based on their underlying ontology structure. Representative GO terms were selected using a similarity threshold of 0.7, and visualizations were generated to highlight major functional categories associated with each gene set.

## Acknowledgments

We extend our thanks to former colony manager Helen Vasquez for her support during the rearing of crowded and isolated specimens. We thank Arianne Cease and Rick Overson at Arizona State University for providing seed colonies of *S. gregaria*. We are grateful for discussions and feedback on the results from multiple members of the Behavioral Plasticity Research Institute research members and in particular, Chuck Zong. Genome assembly was performed in collaboration with the DNA Zoo Consortium (https://www.dnazoo.org), which acknowledges support from Illumina, IBM and Pawsey Supercomputing Center. All opinions expressed in this paper are the authors’ and do not necessarily reflect the policies and views of USDA. Mention of trade names or commercial products in this publication is solely for the purpose of providing specific information and does not imply recommendation or endorsement by the U.S. Government. USDA is an equal opportunity provider and employer.

## Author contributions

Conceptualization: OD, AKC, HAD, SR, FG, GAS, ELA, HS.

Methodology: MAT, SBS, OD, AKC, BF, DMB, RK, DW, SMG, RM, MEP, BS, TJS, ARS, KP, JA.

Investigation: MAT, SBS, OD, EB, DMB, DB, BF, AMCM, MP, SR, ARS.

Visualization: MAT, SBS, EB, DB, AMCM, SR, BF.

Supervision: OD, AKC, SMG, MP, SR, FG, GAS, ELA, HS.

Writing-original draft: all authors.

Writing-review & editing: all authors.

## Data, code and materials availability

All genomic short and long-reads, as well as assembly data generated in this study, have been submitted to the NCBI umbrella BioProjects under accession numbers PRJNA772266 (*S. piceifrons, S. cancellata, S. americana, S. serialis cubense,* and *S. nitens*) and PRJNA814718 (*S. gregaria*). All pairwise genome alignments between species are available on the NCBI Comparative Genomics Viewer at https://www.ncbi.nlm.nih.gov/cgv/. Density-responsive transcriptomic datasets for *S. cancellata* and *S. gregaria* are available through the BioProjects accessions PRJNA798078 and PRJNA1449820, respectively. RNAi transcriptomes of *S. gregaria* are available under the BioProject PRJNA1449934. All scripts and code used for comparative genomics, transcriptomics, and functional validation analysis in this paper are available online at GitHub: https://maevatecher.github.io/locust-comparative-genomics/ (*57*). Raw behavioral assay measurements, OrthoFinder outputs, gene and transcript counts per species, and DEG results are freely available in the GitHub repository for reproducibility of the workflow. Interactive Hi-C contact maps for each species are available on DNA Zoo (https://www.dnazoo.org/).

## RRIDs

RepeatModeler – RRID:SCR_015027

RepeatMasker – RRID:SCR_012954

MCL – RRID:SCR_024109

MAFFT – RRID:SCR_011811

DIAMOND – RRID:SCR_016071

OrthoFinder – RRID:SCR_017118

HyPhy – RRID:SCR_016162

