## Supplementary Material for "Multiple roads to swarming: divergent molecular machineries drive the repeated evolution of locusts"

For papers with only two authors: Paste the full name of the first and second author.

Maéva A. Techer* *et al.*

**This PDF file includes:**

Supplementary Text

Figs. S1 to S11

Tables S1 to S2

**Other Supplementary Materials for this manuscript include the following:**

Data S1 to S09

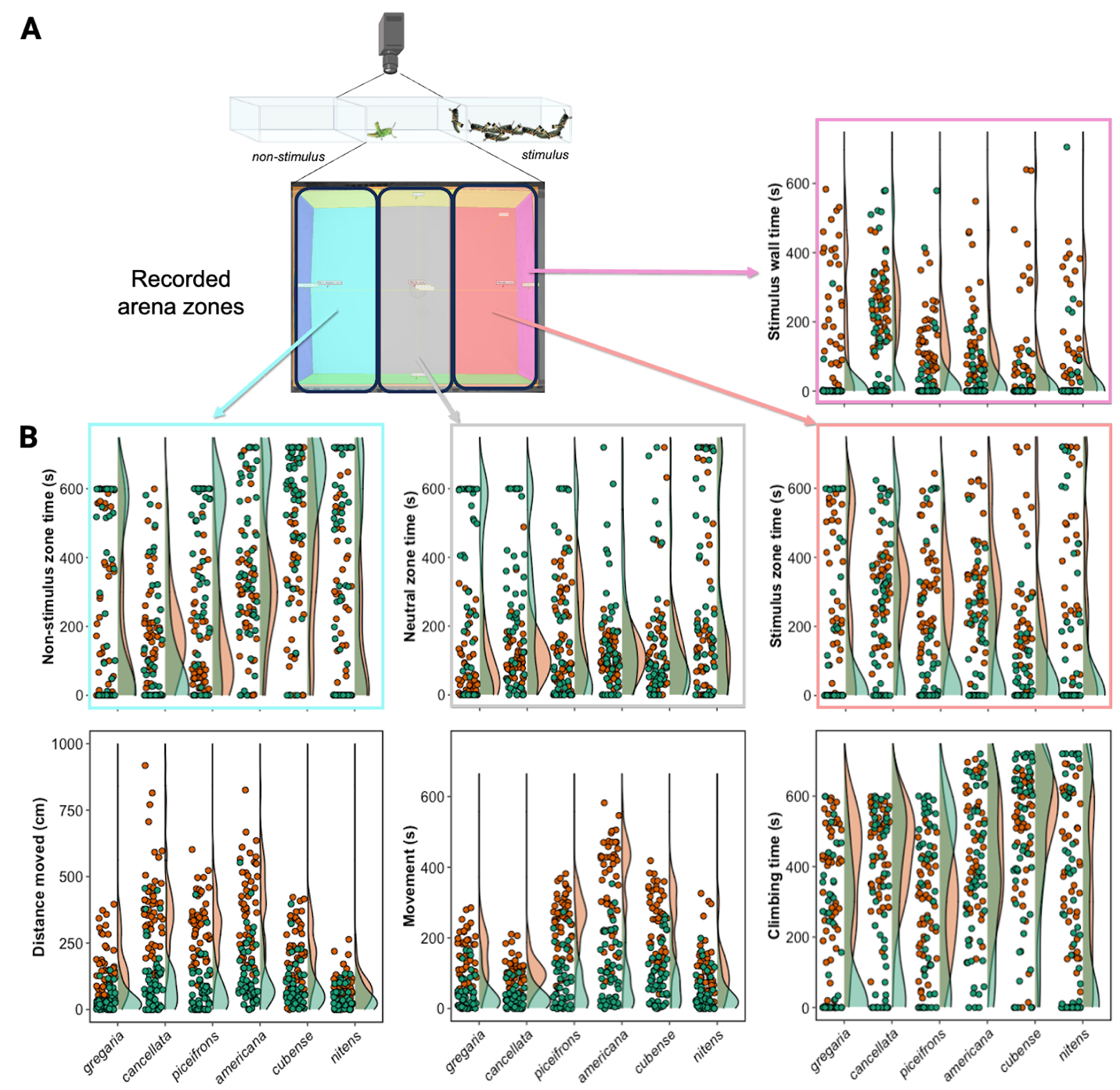

**Fig. S1.**

**Behavioral assays were recorded and standardized in three-chamber arenas.** Panel (A) highlights the recorded arena zones where the focal individual behavior was measured. Panel (B) presents the distribution of the measured behavioral variables, with each point representing a single individual and color-coded by rearing condition (isolated, green; crowded, orange). Measured variables were obtained from 50 biological, independent replicates per condition per species.

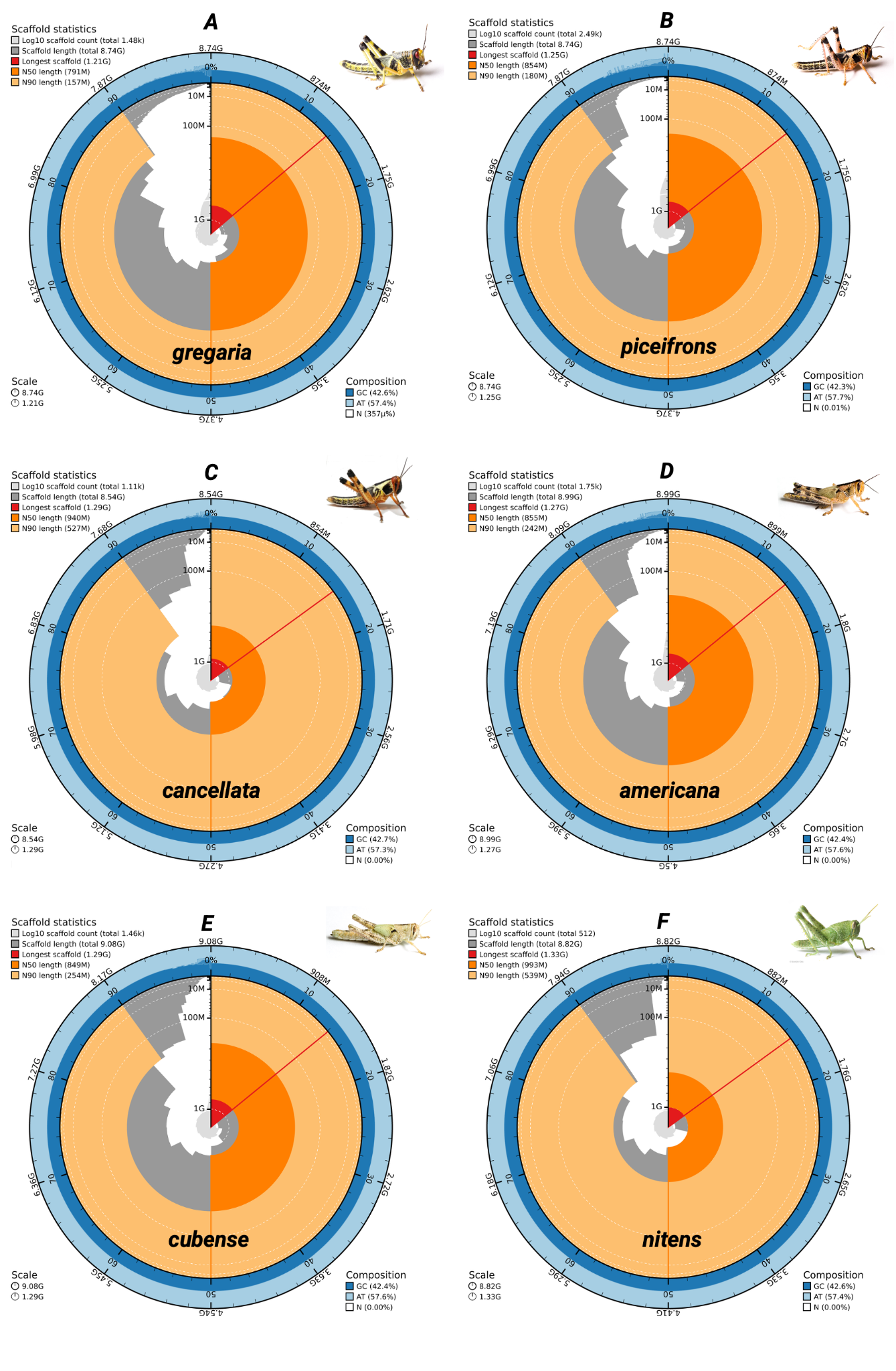

**Fig. S2.**

**Snail plots summarizing genomic metrics of each of the *de novo* assembled *Schistocerca* genomes.** For each assembly, the main plot shows the size of the longest scaffold/chromosome (in red) and the N50 (dark orange) and N90 (light orange) lengths. The dark and light blue rings represent the GC and AT content in the genome, respectively. A picture of a last instar nymph of each species is depicted in the top-right corner of each snail plot. Plots were generated using BlobToolKit and are scaled to each genome size, as indicated in the lower-left corner of each plot. Panels represent A: *S. gregaria*, B: *S. piceifrons*, C: S. *cancellata*, D: S. *americana*, E: S. *serialis cubense,* and F: S. *nitens*.

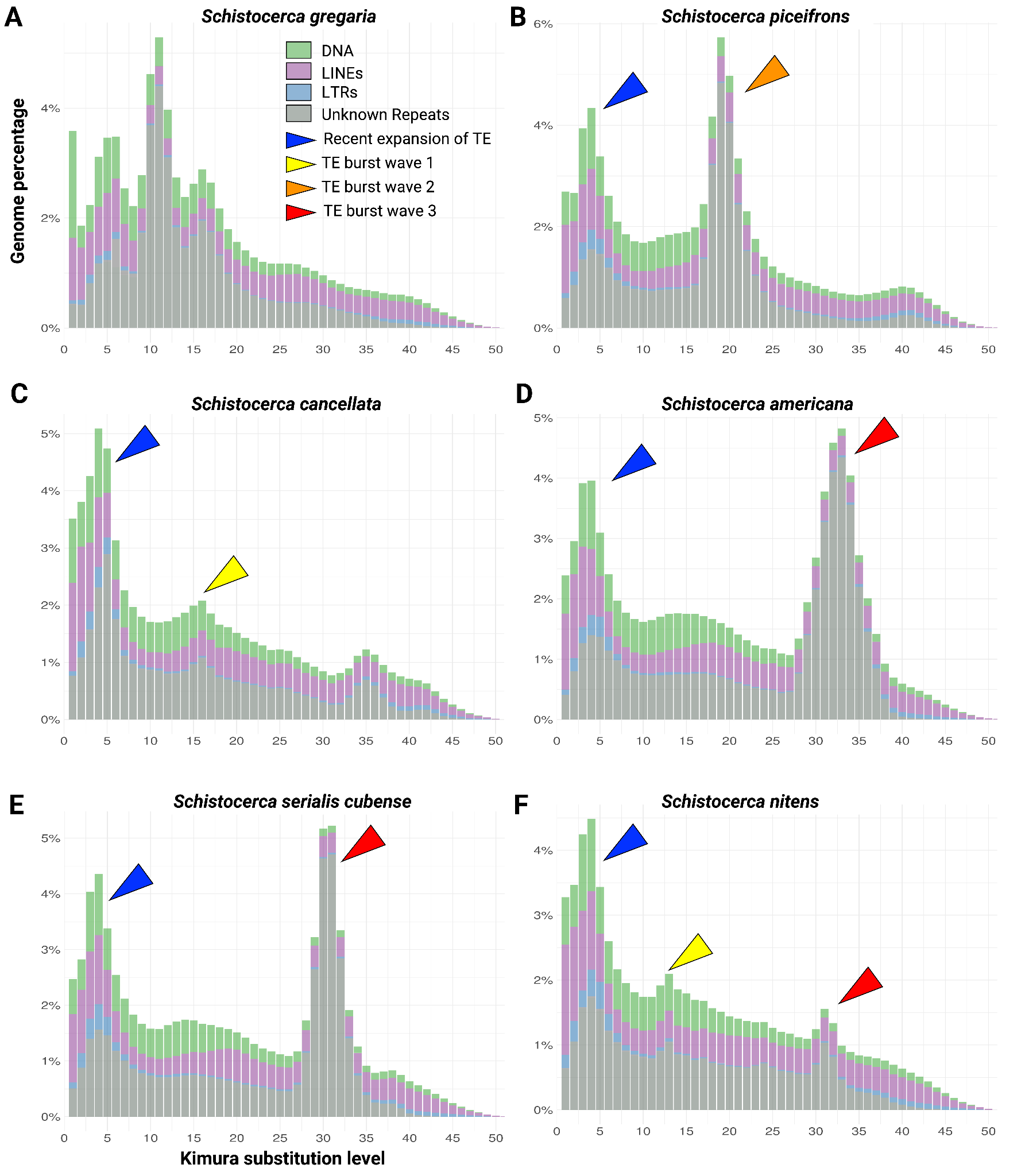

**Fig. S3.**

**Repeat landscape plots of six *Schistocerca* genomes illustrate different histories of transposable element classes expansion and degeneration.** The horizontal axis represents Kimura substitution levels of repeat elements (DNA transposons, LINEs, LTRs, and unclassified repeats) relative to their consensus sequences, serving as a proxy for insertion age: older repeats are positioned to the right, and more recent insertions to the left. The vertical axis shows the genome coverage of each repeat family (in Mb). Colored arrows highlight distinct waves of TE activity: blue arrows mark recent bursts (low divergence), yellow and orange arrows indicate intermediate bursts, and red arrows represent ancient bursts (high divergence, degenerated repeats). Panels represent: (A) *S.* *gregaria*, (B) *S. piceifrons*, (C) *S.* *cancellata*, (D) *S. americana*, (E) *S. serialis cubense*, and (F) *S. nitens*.

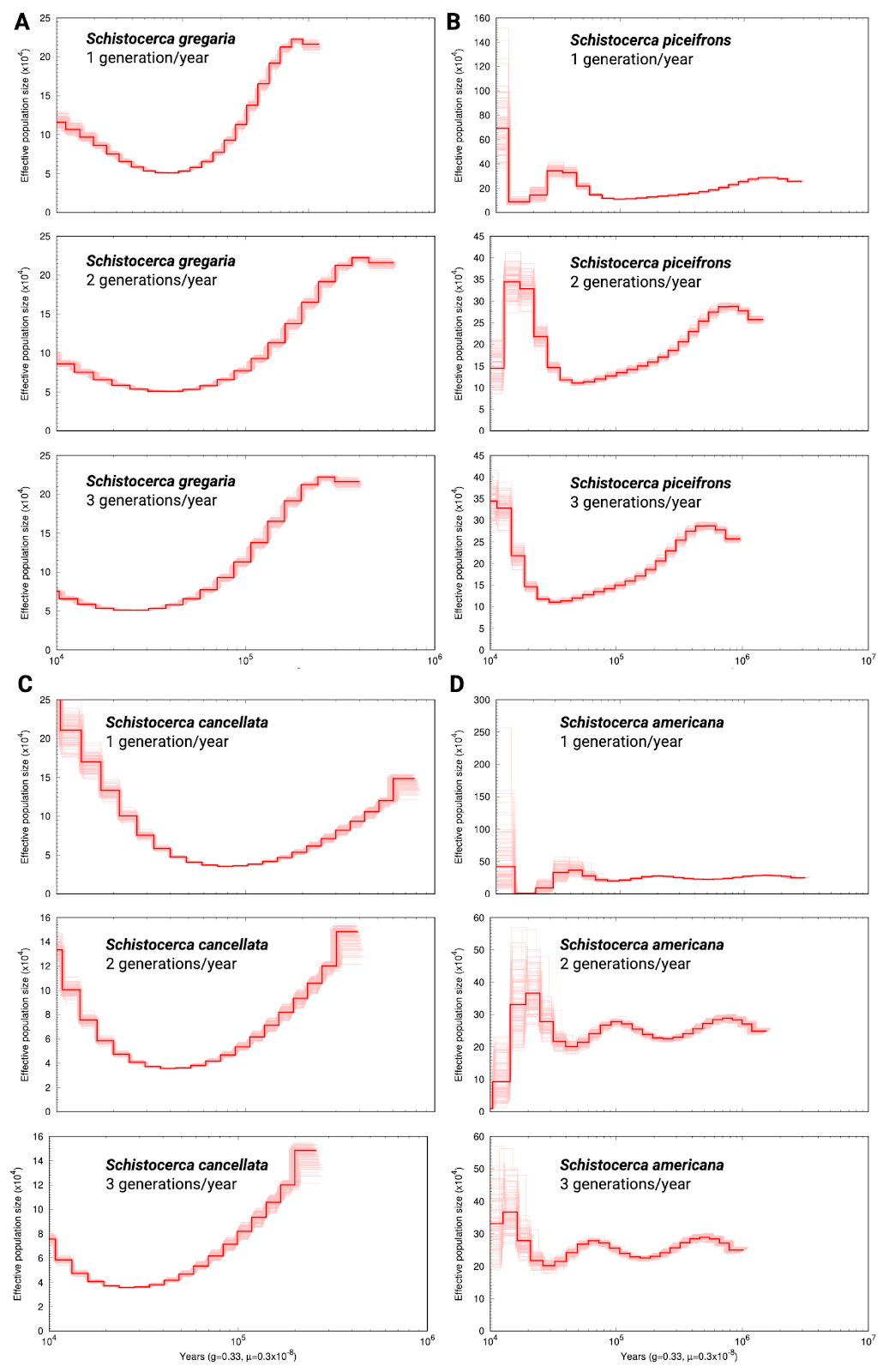

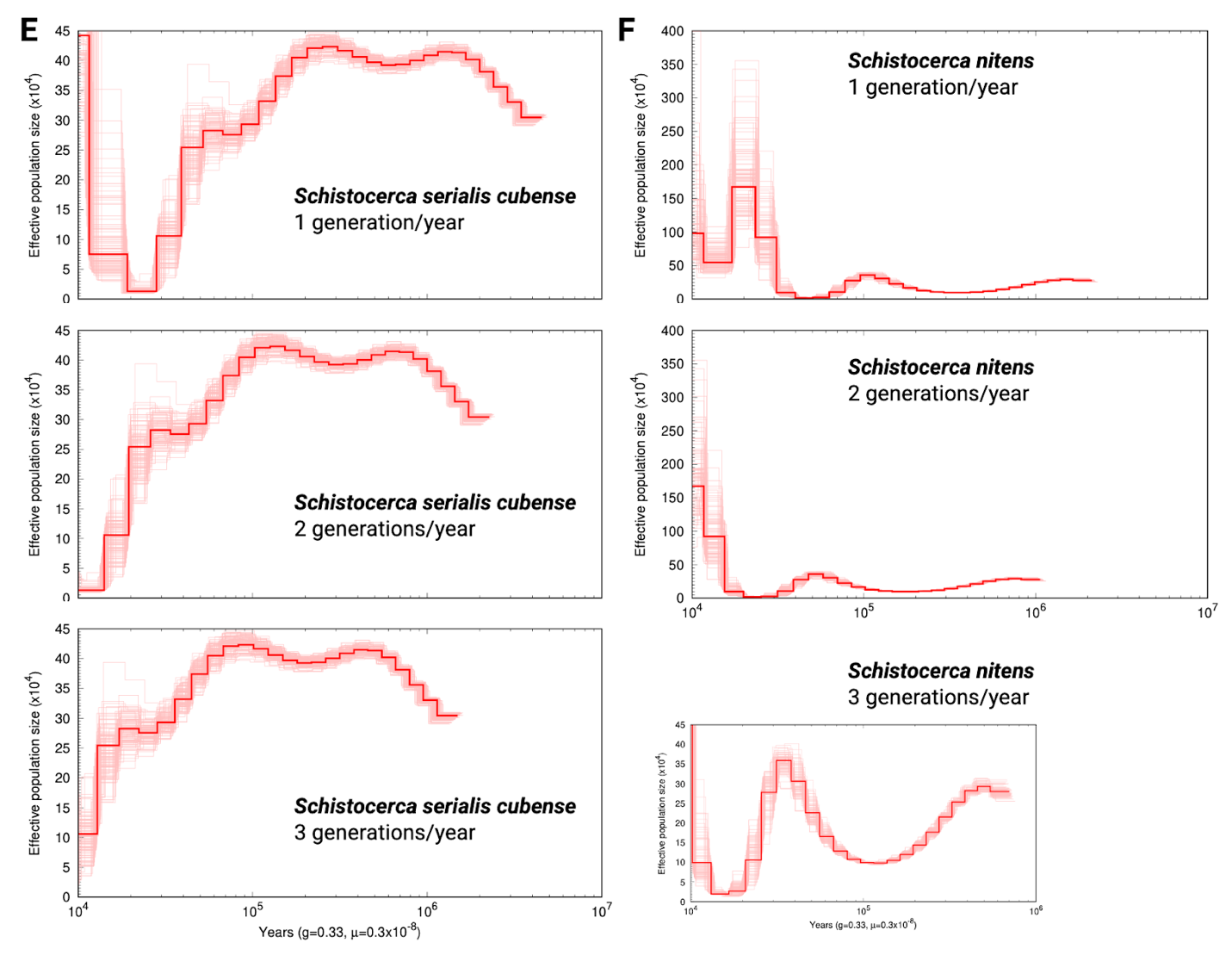

**Fig. S4.**

**PSMC demographic inference plots with various generations per year (from one to three) used for each *Schistocerca* species with *S. gregaria* (A), *S. piceifrons* (B), *S. cancellata* (C), *S. americana* (D), *S. serialis cubense* (E), and *S. nitens* (F).** All panels share the same x-axis, shown as log₁₀-transformed years, and the y-axis indicates the effective population size. Thick red lines indicate the consensus bootstrap model, while the lighter red lines display the confidence interval from the 100 bootstrap runs.

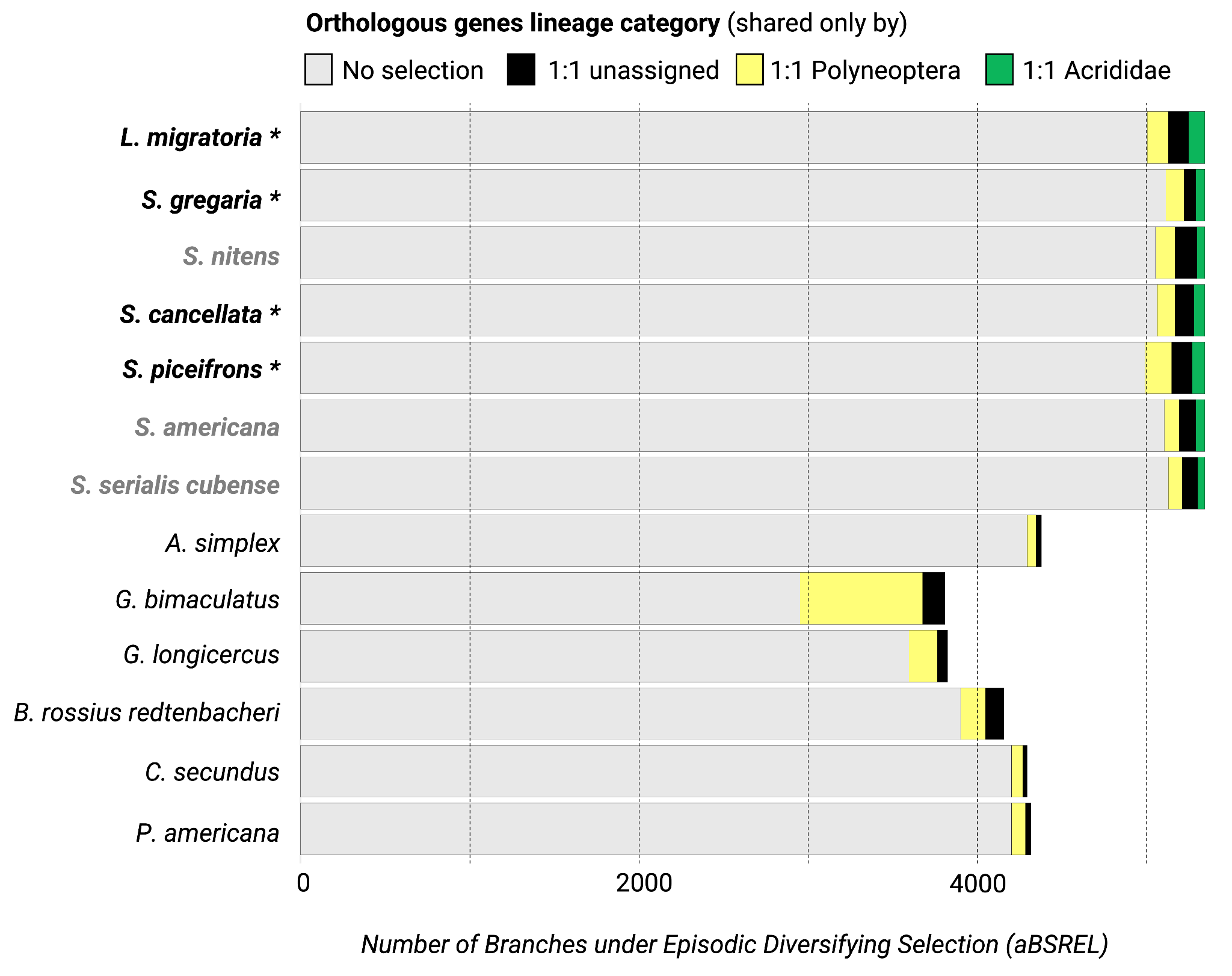

**Fig. S5.**

**Summary results for the exploratory branch episodic positive selection test with aBSREL implemented in HYPHY on single-copy orthologs identified by Orthofinder2.** The results present the number of times the terminal branch (species) had a proportion of sites for each orthgroup that have evolved under positive selection. We classified orthrogroups as Polyneoptera when they were shared (as they were shared by Acrididae and at least one outgroup species), as Acrididae (as they were shared by only *Schistocerca* and/or *Locusta*) or unassigned when we were not sure.
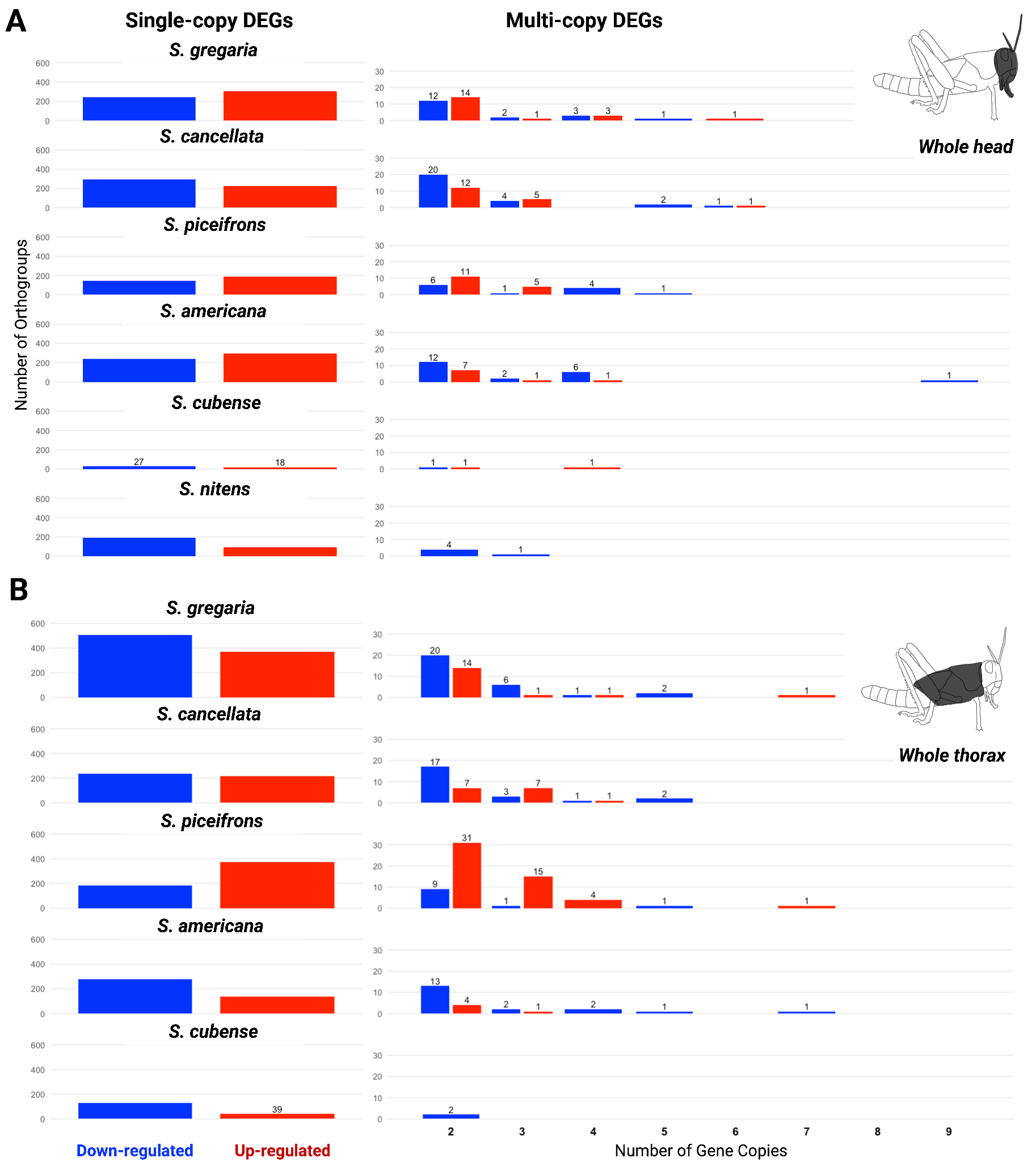

##### Fig. S6.

##### Orthology copy number stratifies density-responsive expression across *Schistocerca* species. We classified density-dependent differentially expressed genes by orthologous copy number in the head and thorax tissues of isolated and crowded-reared female nymphs for each *Schistocerca* species. Directional expression shifts were conserved in trend but heterogeneous in composition: swarming locusts *S. gregaria, cancellata,* and *piceifrons* showed large, tissue-specific DR-DEG sets dominated by copy-variable and multi-copy orthologs, whereas non-swarming relatives *S. americana, serialis cubense,* and *nitens* exhibited progressively attenuated networks and DEG counts.

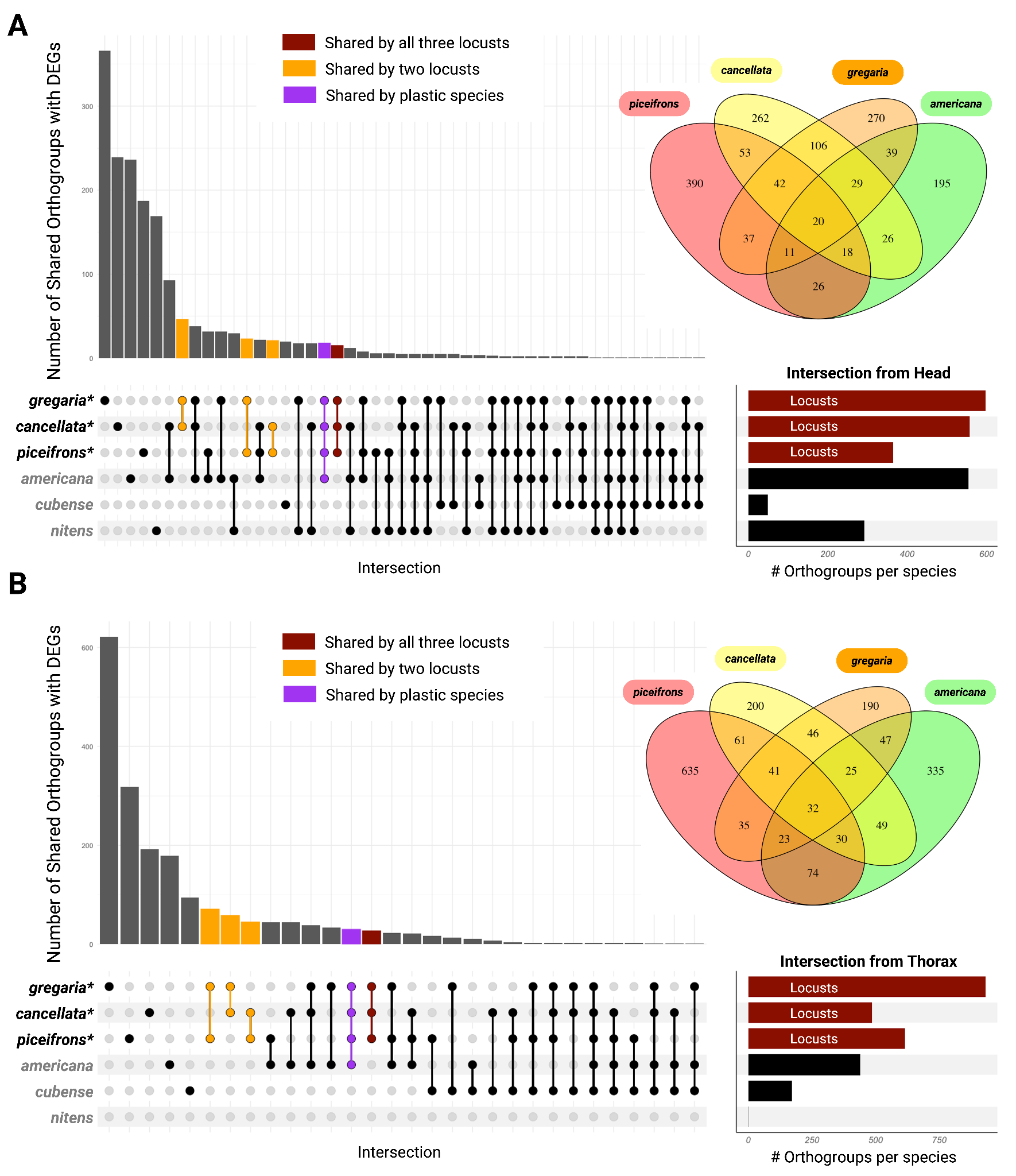

##### Fig. S7.

##### UpSetR plots that summarize cross-species DR-DEG overlaps at the orthogroup level for the head and thorax among all *Schistocerca* species.

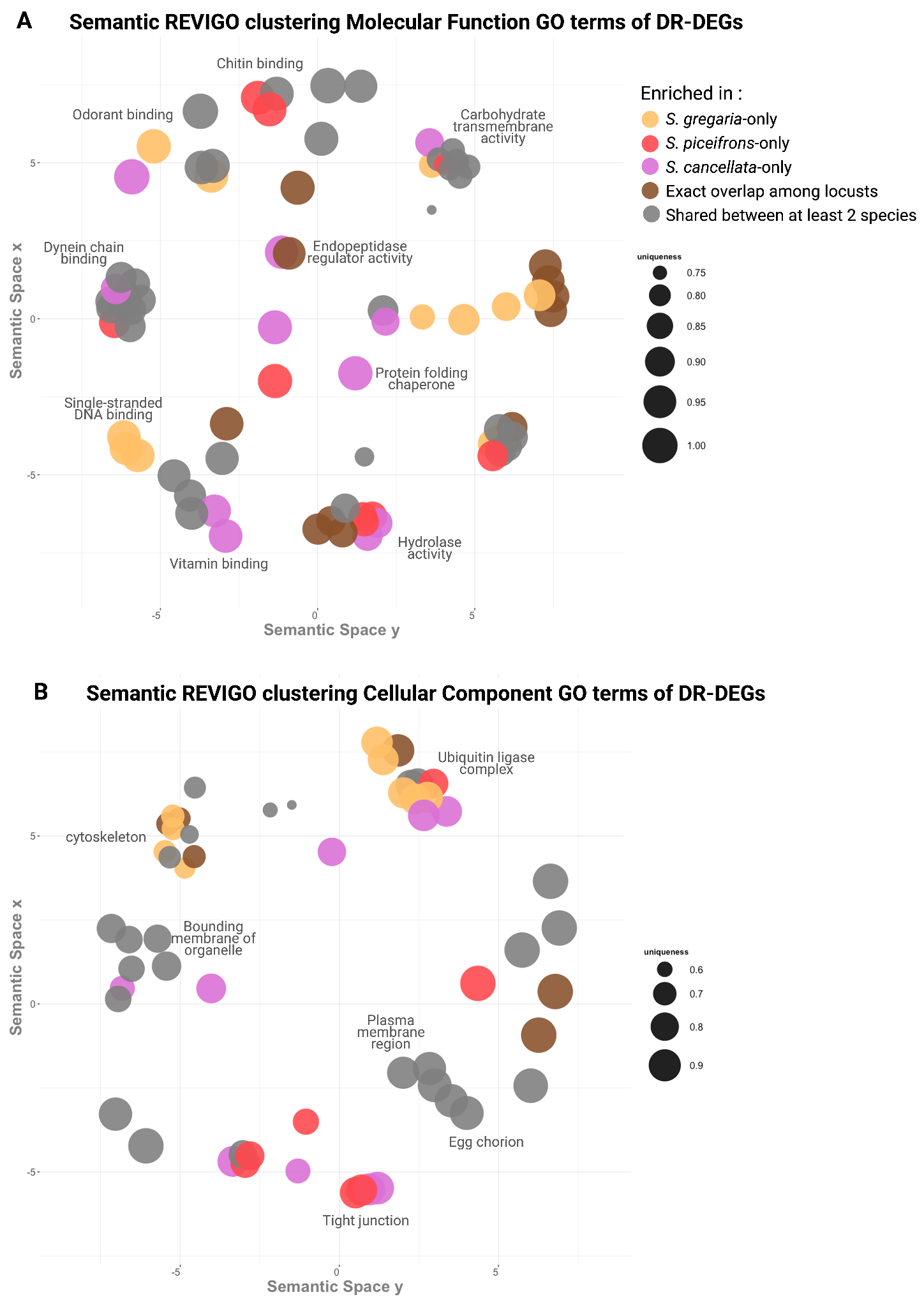

##### Fig. S8.

##### Semantic similarity clustering of molecular function (MF) and cellular component (CC) GO terms enriched among DR-DEGs in swarming locusts. Each circle represents a GO term, sized by the number of genes it contains, with color indicating whether the term is species-specific or shared.

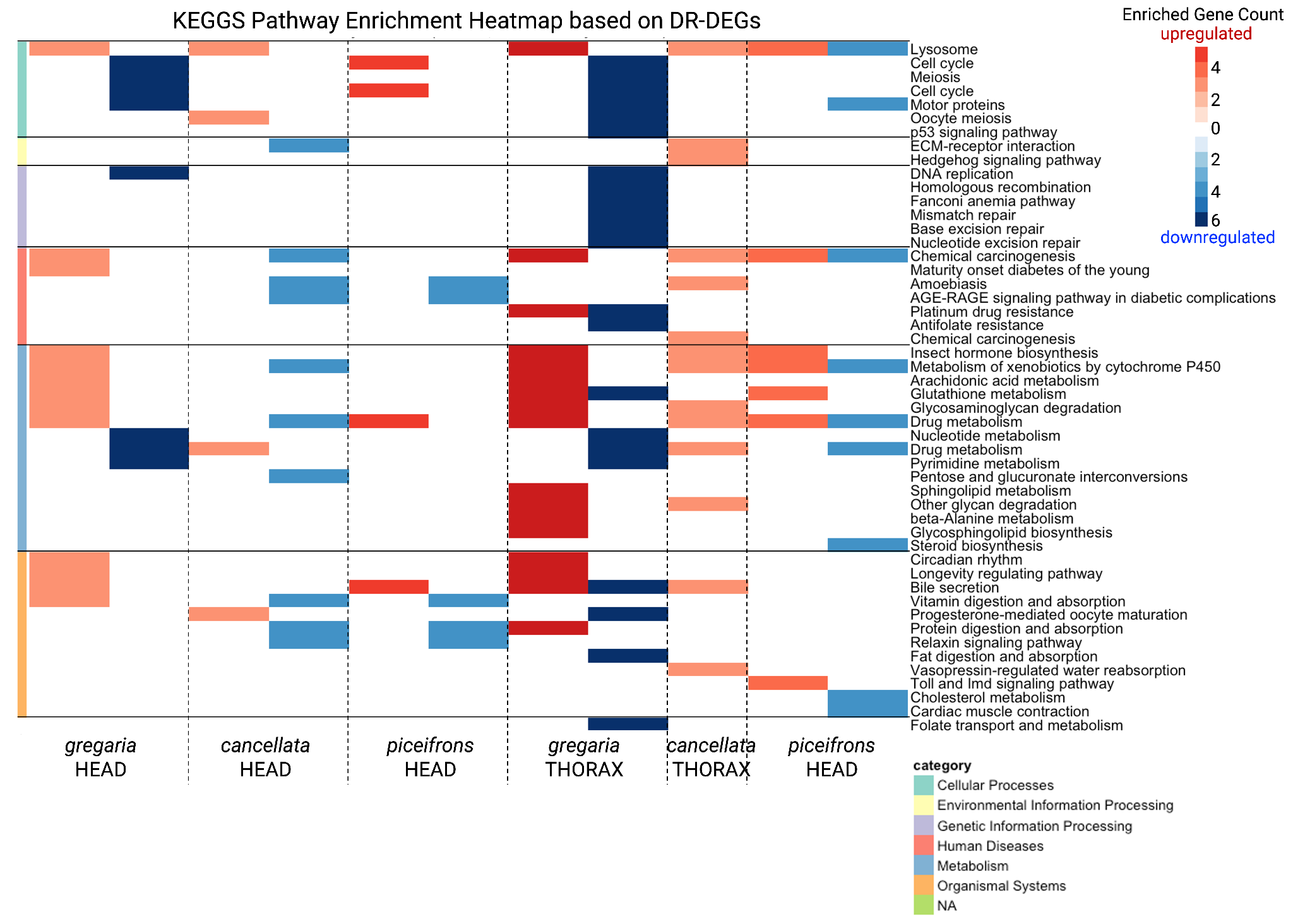

##### Fig. S9.

##### KEGG pathway enrichment heatmap based on the DR-DEGs observed across *Schistocerca* locusts (*S. gregaria,* *S. cancellata,* and *S. piceifrons*)*.*

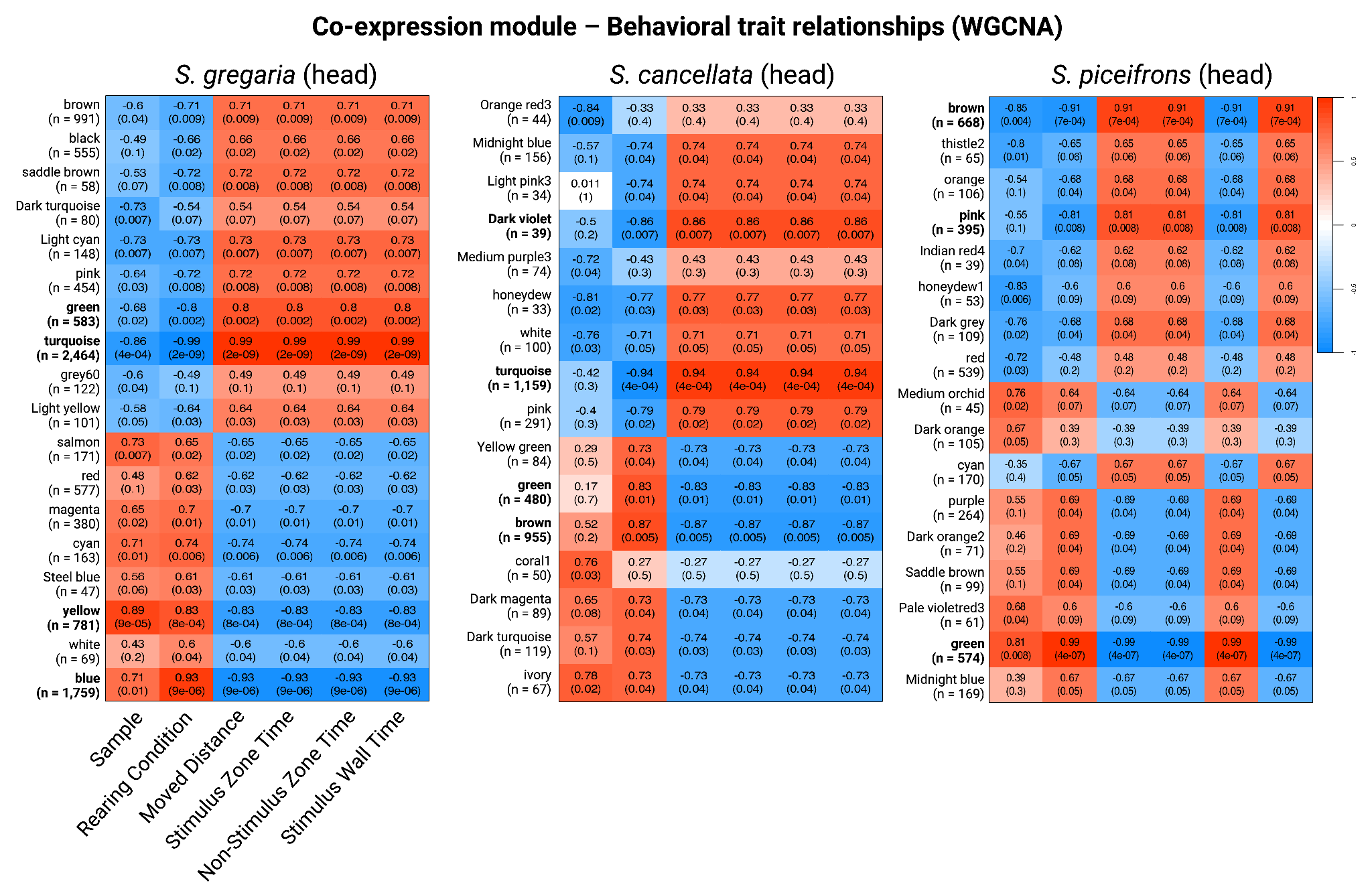

##### Fig. S10.

##### Co-expression module of genes in correlation to behavioral trait in response to a change in rearing density for the three *Schistocerca* locusts (head tissue only). Each correlation matrix represent the significantly filtered gene modules association to different traits (Rearing condition = isolated/crowded, the average total moved distance (cm) = isolated/crowded, the total time spent in the stimulus zone (s), the total time spent in the non-stimulus zone (s), and the total time spent on the stimulus wall (s) averaged for each species and rearing condition). The y-axis shows the name and number of genes in each inferred module, and the color indicates the strength of the correlation. In each case, the number on top indicates the correlation ratio and the number below the p-value.

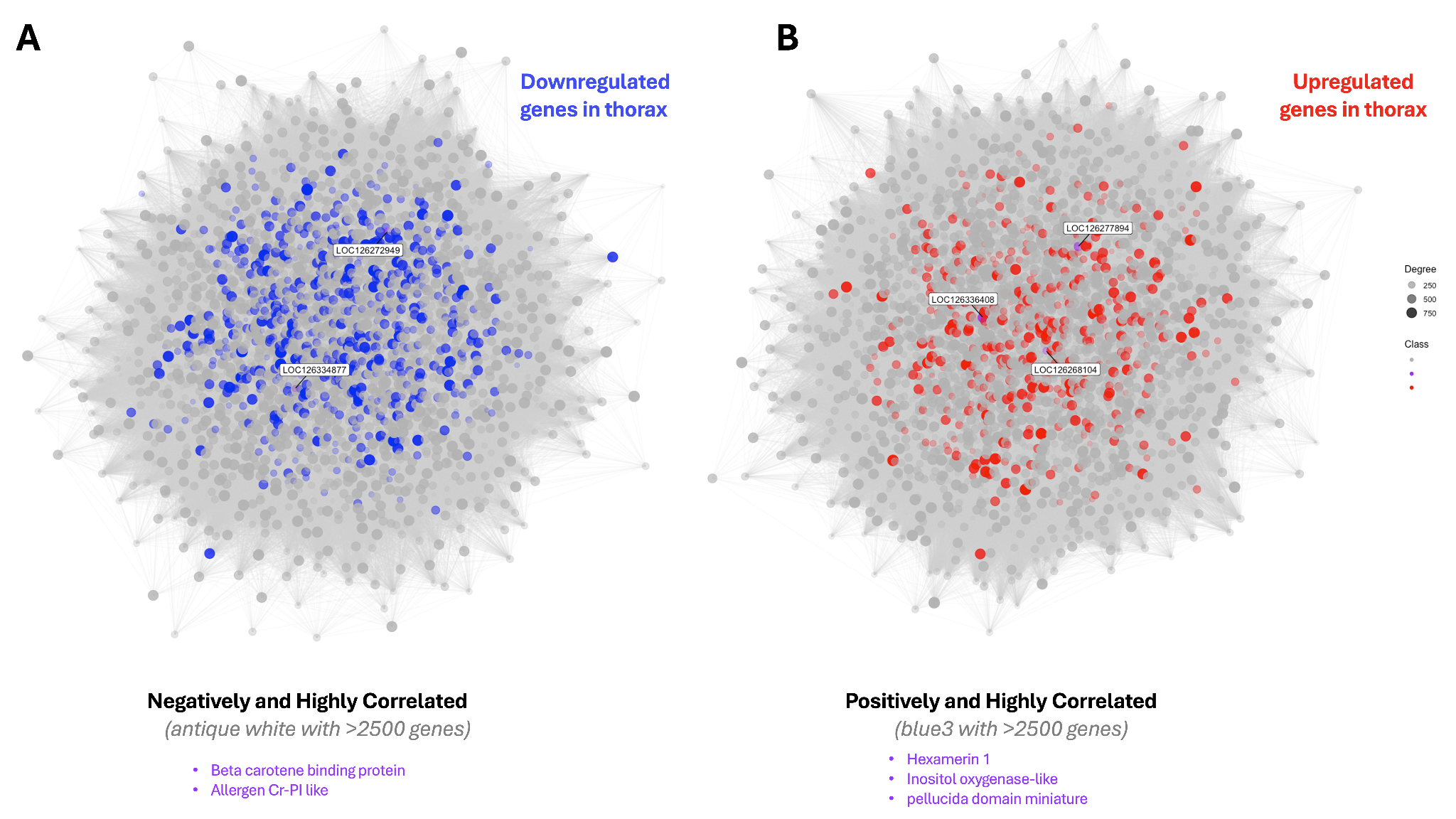

##### Fig. S11.

##### Co-expressed modules strongly associated with behavioral traits (such as total distance moved and time spent in the stimulus zone in the arena) are large and include multiple candidate genes for RNAi in *S. gregaria*. Here, the module network shows each gene and its connectivity ratio via edges (lines) for desert locust thoraxes, based on transcription *vst* values and the median of behavioral trait values, as implemented in WGCNA. Panel A shows a large “antiquewhite” module with downregulated genes, and panel B shows a large “blue3” coded module with up-regulated density-responsive genes. Potential candidate gene positions are highlighted in each network with labels, and their description are indicated below in a purple legend.

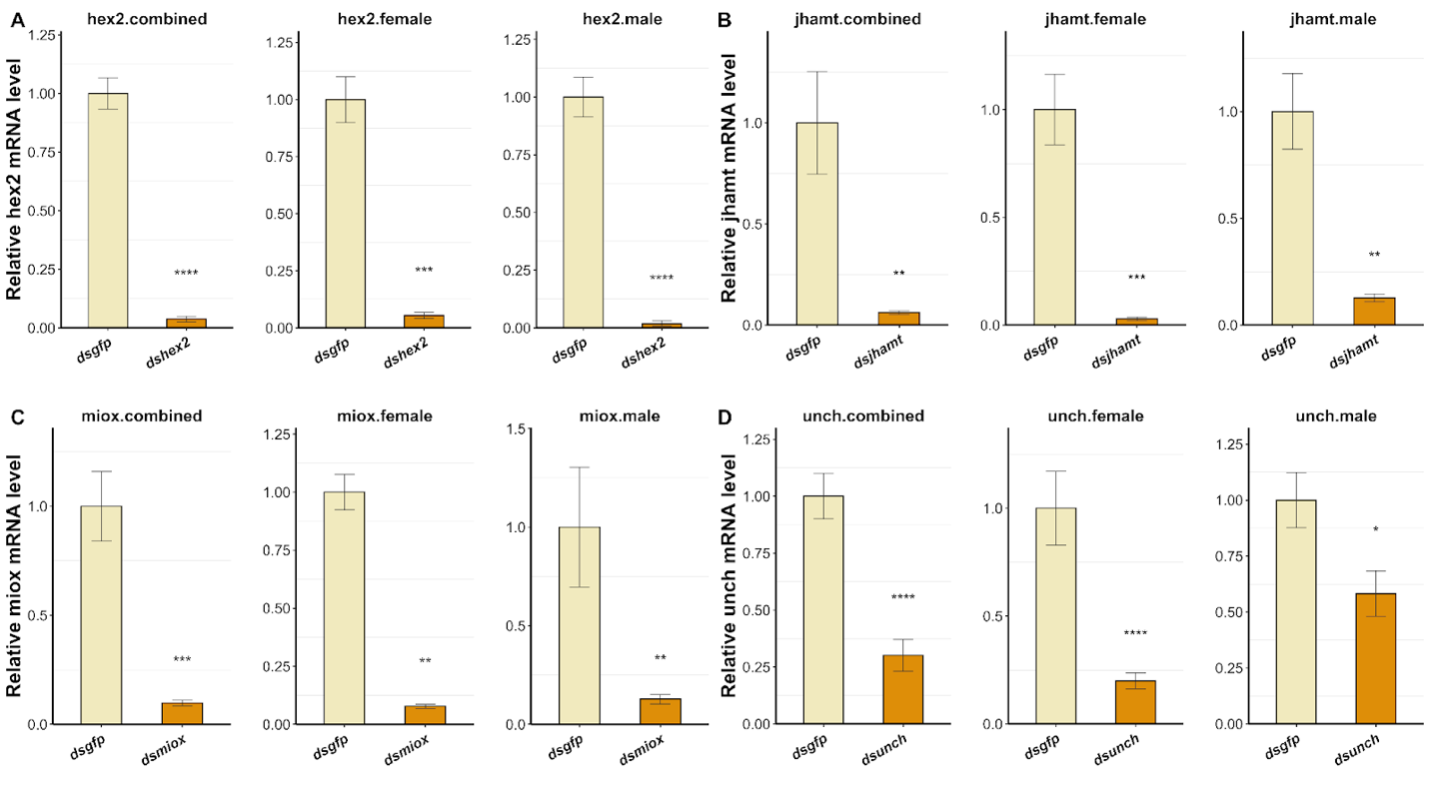

##### Fig. S12.

##### RNAi efficiency in females, males, and combined samples for the targeted genes. (A) *hex2*, (B) *jhamt*, (C) *miox*, (D) *unch*/*m*. Significant differences between the control (*gfp*) and target gene are indicated by asterisks: p<0.05 (*), p<0.01 (**), p<0.001 (***), and p<0.0001 (****).

**Table S1.**

Statistics of the chromosome length assembly generated using HiFi and Hi-C sequencing.

|  | ***S. gregaria*** | ***S. americana*** | ***S. piceifrons*** | ***S. serialis***  ***cubense*** | ***S. cancellata*** | ***S. nitens*** |
| --- | --- | --- | --- | --- | --- | --- |
| **PacBio Sequencing** | 18 cells  326.3 Gb data  37x coverage | 12 cells  300.4 Gb data  34x coverage | 14 cells  242.7 Gb data  28x coverage | 12 cells  251.1 Gb data  28x coverage | 12 cells  248.2 Gb data  29x coverage | 17 cells  270.6 Gb data  31x coverage |
| **Total length** | 8,742,434,016 | 8,990,369,808 | 8,742,459,392 | 9,082,822,131 | 8,536,893,415 | 8,822,571,301 |
| **Scaffold count** | 1,475 | 1,751 | 2,493 | 1,455 | 1,108 | 512 |
| **Scaffold N50** | 791,150,472 | 854,857,182 | 854,032,331 | 849,285,246 | 939,673,033 | 992,907,699 |
| **Contig count** | 1,787 | 2,248 | 3,413 | 1,968 | 1,511 | 827 |
| **Contig N50** | 4,4584,537 | 49,627,568 | 27,424,730 | 39,994,732 | 59,628,095 | 73,929,931 |
| **Total gap length** | 31,200 | 248,500 | 460,000 | 255,132 | 197,448 | 157,037 |
| **GC %** | 42.5 | 42.5 | 42.5 | 42.5 | 42.5 | 42.5 |
| **Protein-coding** | 19,799 | 17,662 | 17,490 | 17,237 | 16,907 | 17,500 |
| **non-coding** | 75,804 | 56,384 | 69,257 | 51,178 | 80,053 | 48,771 |
| **Pseudogenes** | 3,862 | 7,227 | 10,058 | 7,394 | 6,571 | 6,288 |
| **BUSCO score %** | 99.2 | 99.4 | 99.3 | 99.3 | 99.3 | 99.3 |

##### Table S2.

##### Primers used for functional validation with RNAi.

| Gene Name | Sequence (5’-3’) |
| --- | --- |
| dshex1 | F: TAATACGACTCACTATAGGGTGCTCCGTGACCCATTGTAC  R: TAATACGACTCACTATAGGGCCTCATCCACCGTCATCTCG |
| dshex2 | F: TAATACGACTCACTATAGGGTCATTCACTGTCGCTGTGCT  R: TAATACGACTCACTATAGGGCAGGCAAGTGGTTGGAAAGC |
| dsjhamt | F: TAATACGACTCACTATAGGGAGCATGCACCTTACGGGAAA  R: TAATACGACTCACTATAGGGAGGTTGGCTTCCCATACTGC |
| dsmiox | F: TAATACGACTCACTATAGGGGAACAAGCTGGTGGACGAGA  R: TAATACGACTCACTATAGGGGCGCTATTTCTGGCAGCTTC |
| dsunch/dsm | F: TAATACGACTCACTATAGGGGAACAGCCACATCCTGGACA  R: TAATACGACTCACTATAGGGGCACCTGGTACCGGTATAGC |
| hex2 qPCR | F: CCGTGTCTTCATTGGTCCCA  R: TCTCACCAACGGACAGCTTC |
| jhamt qPCR | F: GAAGCACAACGCACTTCCAG  R: CTGCTCCAGGTTGCGAGATG |
| miox qPCR | F: GACGAGACGCCAGACATTGA  R: GCTTCCAGGCAGGTACTTGT |
| unch/m qPCR | F: GACCTTCTTCTCGTTCCGCA  R: CTTCGCTTCATCCCACACCT |

**Data S1. (separate file)**

Summary and details of repetitive elements annotated for each *Schistocerca* genome.

**Data S2. (separate file)**

List of orthogroups (gene family) inferred from Polyneoptera genomes and protein phylogeny using OrthoFinder2 (inflation 2).

**Data S3. (separate file)**

List of orthogroups (gene family), gene count, and clade assignment.

**Data S4. (separate file)**

Parsed aBSREL results implemented on HYPHY, on single-copy orthogroups inferred from OrthoFinder2 on the Polyneoptera phylogeny using *Schistocerca* genomes.

**Data S5. (separate file)**

Parsed BUSTED-PH and RELAX results implemented on HYPHY, on single-copy orthogroups inferred from OrthoFinder2 on the Acrididae pruned and whole phylogeny trees.

**Data S6. (separate file)**

Read mapping statistics to the own RefSeq versus the shared *S. gregaria* RefSeq using STAR-2 passes mode.

**Data S7. (separate file)**

Significant differential gene expression statistics, following surrogate variable analysis correction on DESeq2.

**Data S8. (separate file)**

List of genes of interest according to locust biology and/or Density Responsive-DEGs analysis. Candidate genes selected for probe design, knockdown efficiency test, and final functional RNAi are highlighted in yellow.

**Data S9. (separate file)**

Metadata and sequencing statistics from whole transcriptome bulk tissues in *Schistocerca* species
